# The comparative strengths and limitations of Nile Red and 9-(dicyanovinyl)-julolidine (DCVJ) fluorescent dyes for detecting microplastics and nanoplastics

**DOI:** 10.64898/2026.08.03.742549

**Authors:** Madison Wallner, Justin Diaz, Amelia B. Labbe, James J. Jacob, Quentin Williams, Adina Paytan, Clive R. Bagshaw

## Abstract

Nile Red is widely used for the detection of microplastics because its fluorescence emission is sensitive to local polarity and can distinguish hydrophobic plastics from hydrophilic ones. The fluorescence of the molecular rotor, 9-(dicyanovinyl)-julolidine (DCVJ) is less sensitive to polarity but more to viscosity. DCVJ is less widely used for microplastic analysis, although it has been used to detect polystyrene nanobeads. Here, we compared these dyes with standard samples from the Hawai’i Pacific University Polymer Kit 1.0 and confirmed that Nile Red, in general, was better for the detection and identification of microplastics. Fluorescence emission was analyzed using photography, as well as spectroscopy. The color and peak emission wavelength of some stained environmental microplastics were affected by additives. Raman spectroscopy was used to confirm the chemical identity of such samples. Although DCVJ emits green fluorescence on binding to some microplastics, a peak at 620 nm has been reported with polystyrene nanobeads, attributed to dimer/excimer formation. We confirmed this property and directly observed diffraction-limited spots using fluorescence microscopy, attributed to single or just a few nanobeads. Nile Red also stains polystyrene nanobeads and gave stronger signals than with DCVJ, but Nile Red was prone to false positives due to dye aggregation in aqueous solutions.

## Introduction

Research has shown that microplastic pollution is detectable in all regions on Earth. Microplastics have been detected in the ocean [1,2], freshwater systems [3,4], aquatic sediments [5], terrestrial environments [6–8], and even remote areas [9]. While there is variation in the stated size limits, microplastics have been defined as anthropogenic polymeric particles with all three dimensions greater than 1 nm and smaller than 5 mm, to which chemical additives or other substances have been added. Nanoplastics define similar materials but a smaller subset where the stated size limits range from 1 - 100 nm to 1-1,000 nm [10,11]. Thus, nanoplastics are near or below the resolution limit of standard light microscopy and may be small enough to cross cell membranes. Microplastics enter the environment from both primary and secondary sources. Primary sources of microplastics are those that were manufactured in a size class smaller than 5 mm, such as microbeads in face scrubs or raw resin ‘nurdles’. Secondary microplastics are produced from the degradation and breakdown of larger plastic items in the environment [12].

Micro- and nanoplastic pollution poses significant environmental and biological risks due to the combined effects of chemical additives present in plastics and the diverse exposure pathways of polymer particles into organisms. Many additives, such as plasticizers, antioxidants, and flame retardants, are not chemically bound to the polymer matrix and can migrate into surrounding aquatic and terrestrial systems, where they may affect organisms [13]. Some of these substances are classified as endocrine-disrupting chemicals and, upon ingestion, can interfere with behavioral and reproductive functions [14,15]. Other plastic additives have been identified as carcinogenic, immunosuppressive, acutely toxic, and, in certain cases, lethal [15–17]. Micro- and nanoplastics can adversely affect individual organisms through ingestion, as well as exert harmful effects across trophic levels [18]. Microplastics have been detected in multiple locations of the human body, including lung tissue [19], placenta [20], bronchoalveolar lavage fluid [21] and blood [22]. As concentrations of MP particles increase, so do their chronic and adverse health effects on organisms [20].

Given the harmful nature of micro- and nanoplastic contamination, much effort has been directed towards detection and identification methods. Fluorescence microscopy is widely used for the detection of microplastics, particularly in conjunction with lipophilic stains such as Nile Red [23–27]. This method has the advantage of being sensitive and can be carried out with relatively simple instrumentation. The solvatochromatic properties of Nile Red emission potentially allow the broad identification of plastic types in terms of hydrophobicity but the measurement cannot identify specific chemical structures, unlike infrared and Raman spectroscopy. Other limitations include the problem of false positives from the staining naturally occurring hydrophobic materials (e.g. lipids) and false negatives due to strongly light-absorbing pigments present in some plastics that quench the dye fluorescence. Ho et al. [27] has critically reviewed the potential for fluorescence staining as a stand-alone method for identifying microplastics. However, even with its limitations, fluorescence detection remains a useful way to monitor microplastics in two scenarios. When combined with techniques such as infrared and Raman spectroscopy, it provides a way of pre-screening the sample for candidate microplastics, so increasing the workflow [28]. Secondly, the use of simple instrumentation makes the approach amenable to citizen scientists, as well as school and college students [29–31], thus increasing awareness and the geographical areas that can be sampled for microplastics [32].

Nile Red remains the most widely used dye to stain microplastics, but alternatives have been explored. In particular, the fluorescent molecular rotor, 9- (dicyanovinyl)-julolidine (DCVJ), has been shown to stain polystyrene spheres with dimensions of 50 to 100 nm, thus extending the methodology to the nanoplastic scale [33–35]. Molecular rotors show enhanced emission on binding to hydrophobic surfaces by restricting the rotation around a single C-C bond and so reducing the probability of radiation-less decay back to the ground state. DCVJ is commonly used to monitor local viscosity within cells, but its application to microplastic detection has been limited [36]. Here, we compare the efficacy of Nile Red and DCVJ (Fig 1) for detecting microplastics in general.

**Fig 1.**
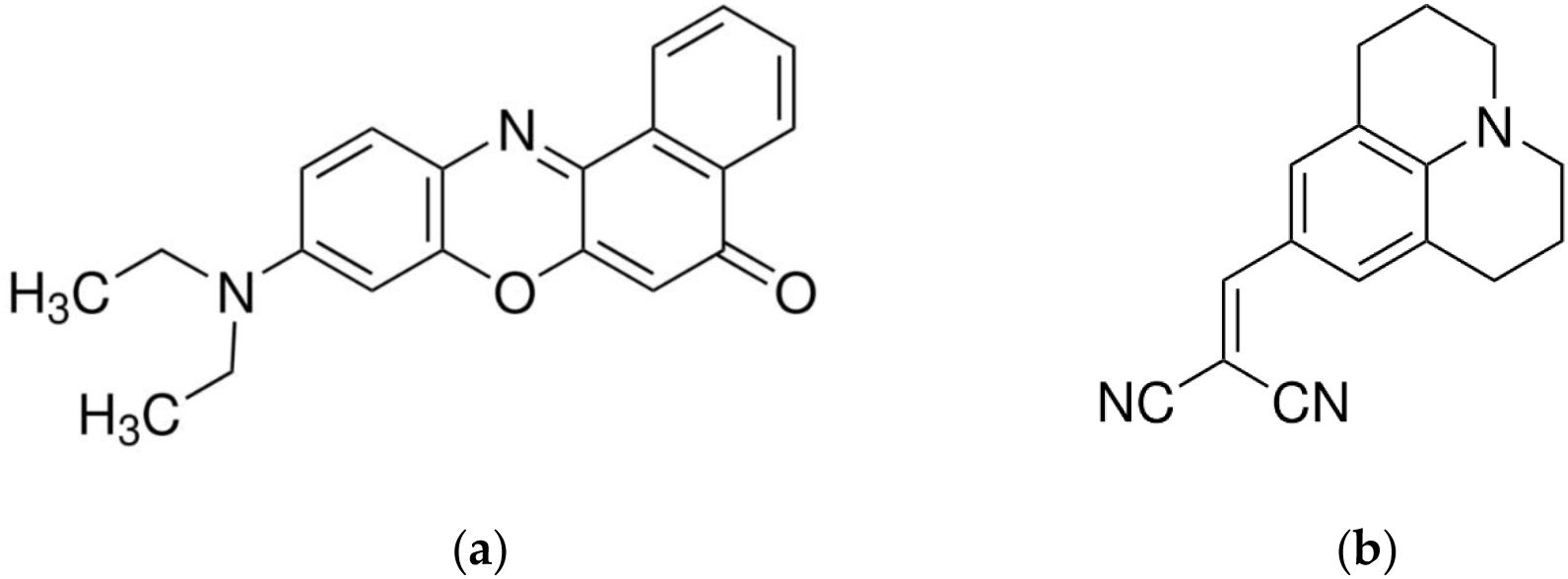
The structures of fluorescent dyes. (a) Nile Red and (b) 9-(dicyanovinyl)- julolidine (DCVJ).

A challenge of using fluorescence detection for microplastics is establishing a standard protocol so that results from different studies can be compared. This is difficult because the degree of staining is sensitive to the experimental conditions used, including solvent composition, temperature and incubation time. Moreover, even with identical staining procedures, the reported fluorescence intensity and color depend on the instrumentation used for detection [37]. Factors such as the wavelength and bandwidth of the excitation source, the spectral sensitivity of the detector and the transmission properties of excitation and emission filters in the optical path all contribute to the observed signal. One approach to dealing with this challenge is to include standard plastic samples. Here, we use the Polymer Kit 1.0 compiled by the Hawai’i Pacific University and the Center for Marine Debris Research [38] to compare staining with Nile Red and DCVJ using photographic and spectroscopic detection.

## Materials and Methods

### Materials

Nile Red was purchased from MP Biochemicals (via VWR Ltd) and 9- (dicyanovinyl)-julolidine (DCVJ) was obtained from Cayman Chemical (Ann Arbor). 70 nm (50-100 nm) Sphero^TM^ polystyrene beads were supplied by Spherotech Inc (Lake Forest). The Polymer Kit 1.0 was obtained from the Hawai’i Pacific University [38] and contained polyethylene (PE) of differing densities (LDPE, MDPE, HDPE), polypropylene (PP), polyethylene terephthalate (PET), ethylene-vinyl acetate (EVA), acrylon-butadiene-styrene (ABS), polystyrene (PS), polyamide (Nylon, PA), polyvinyl chloride (PVC) pellets (∼3 to 5 mm diameter) and polyester fabric (PEST). Other plastics were derived from consumer packaging and environmental samples.

### Staining procedures

Stock solutions of Nile Red and DCVJ were made up at 1 mg/mL in isopropyl alcohol (IPA), and sub stocks made at 0.1 mg/mL. Stock concentrations were checked by absorption using a Thermo Genesys 10S uv/vis spectrophotometer (Nile Red in DMSO, E_554_ = 19,600 M^-1^cm^-1^, [39]; DCVJ in methanol, E_454_ = 62,000 M^-1^cm^-1^, [40]). Staining of Polymer Kit samples was performed at a dye concentration of 8 µM (2 µg/mL DCVJ or 2.5 µg/mL Nile Red) in 25% acetone, 75% H_2_O at 35 °C on a shaking hotplate. This acetone-water mixture was deemed optimal for staining a range of plastics [27]. Individual pellets were arranged in a 96 well plate and covered with 100 µL of the dye solution. In addition, control samples were run with solvent only to check for any intrinsic fluorescence from the pellets. After 1 hour, the pellets were rinsed in millipore H_2_O and allowed to dry. The samples were stored in the dark prior to analysis. The pellets were transferred to a black metal tray for photography (Fig S1a) or a modified cuvette for spectrophotometric analysis (Fig S3). Pellets were held in place using double-sided tape which itself showed no significant fluorescence. The same procedure was used to stain environmental microplastics, including those collected from Waimānalo beach, O’ahu, Hawaii.

### Photography

Stained pellets were arranged on a gridded black metal tray and illuminated with a blue (450 nm LED) flashlight (Xinfeibei) without a focusing lens, held at 45° and 15 cm away. Photographs were taken with Nikon D3000 10.2 MP dSLR camera with a 55 mm lens set at f/16 aperture and exposure times 0.25 to 1 s. This camera has 3,872 x 2,592 pixels with an area of 6.1 µm square. A Zeiss 515 nm long-pass filter (Fig S2) was attached to the camera lens using filter stepping rings. The small camera aperture was selected to enable the use of a 25 mm diameter filter and allow for direct comparison with microplastic samples photographed through a microscope with the same filter. Files were saved in JPG and NEF (RAW) format and analyzed using ImageJ [41]. With JPG files, RGB values were determined by selecting a Region of Interest and using the Plugins/Analyze/RGB Measure option or converting the image to 3 channels using Image/Color/Split channels, then selecting a Region of Interest. Raw NEF images were imported as a hyperstack using the Bio-Formats plugin and then converted to a stack using the Image/Stacks/Hyperstack to Stack option to give the three RGB channels in separate frames. A Region of Interest was then selected and RGB values determined using Image/Stacks/Measure Stack.

### Fluorescence spectroscopy

Fluorescence emission spectra were recorded using a Varian Eclipse spectrofluorimeter. Typically, excitation was set at 450 nm and the emission scanned from 470 to 800 nm with 5 nm slit widths and photomultiplier voltage from 600 to 900 V. This excitation wavelength was selected to allow a direct comparison of Nile Red and DCVJ, and also to explore the large range of emission wavelengths (520 nm to 640 nm) observed with Nile Red with minimal contributions from light scattering. Liquid samples and nanosphere polymer suspensions were recorded using a 1 cm or 2 mm pathlength cuvette. Solid film-like samples were mounted in a custom holder made from a Thorlabs 25 mm FMP1 mirror holder mounted on a BA1 base (Fig S4c). The plastic film was held by 5 mm magnets directly or via a coverslip and held by surface tension from 10 µL of H_2_O. Transparent films were angled at 45° to reflect the excitation beam away from the detection port (Fig S4b). Polymer pellets were mounted in a modified acrylic cuvette with 3 sides cut-away and the sample mounted on double-sided tape at a height of 15 mm (Fig S3). The pellet was centered in the 450 nm excitation beam using the height adjustment screw to achieve maximum visual scattering.

The reported emission spectra are uncorrected for the instrument sensitivity which gives a stronger signal in the green compared with the red region. While the correction factors for the Varian fluorimeter were not available across the complete wavelength range of interest, representative values for an SLM fluorimeter having the same R928 photomultiplier are: 500 nm, 1.00; 550 nm, 1.57: 600 nm, 2.64; 650 nm, 5.17; 700nm, 11.9.

### Fluorescence microscopy

Three different microscopes were used to examine plastic samples stained with Nile Red and DCVJ, each having different features.

i. An Amscope 120 microscope was adapted for fluorescence as previously described, using a 450 nm laser pen for illumination [29]. The same Nikon 3000 dSLR camera, as used for photography above, was attached to the trinocular port via a wide aperture 10x EagleEye digiscoping eyepiece. Exposure times ranged from 0.25 to 30 seconds. This set-up comprises consumer-grade equipment which makes it accessible for citizen science and classroom activities. A research-grade Zeiss 515 nm long pass filter was used to block the excitation light to allow direct comparison with the macroscopic images of the standard polymer kit and environmental plastics. However, lower grade acrylic filters can be used for demonstration purposes [29], at the expense of a larger contribution from breakthrough excitation light (Fig S1).
ii. A modular fluorescence microscope was constructed incorporating a 20X Mitutoyo 0.42 NA infinity-corrected, long working-distance microscope objective, in conjunction with a 200 mm focal length tube lens and a 1/1.2” CMOS camera (FLIR Blackfly BFS-U3-23S6C) with a color sensor consisting of 1980 x 1200 5.86 µm square pixels (Sony IMX249). Excitation was provided by a 40 mW 450 nm diode laser, packaged in a 12 mm diameter tube and mounted external to the microscope body, at an angle of 70 degrees to the microscope axis. The laser was held in a kinematic mount to allow for alignment. The beam was brought to near focus at the sample with a 100 mm focal length lens. A Schott OG530 filter (Fig S2a) was used to block excitation light from reaching the camera. Images were typically recorded with a frame rate of 4.2 Hz. The design of this microscope was driven by the long-term goal of combining fluorescence microscopy with the optics required for stimulated Raman microscopy using infrared lasers.
iii. A prism-based, total internal reflection fluorescence (TIRF) microscope, developed for detecting single fluorophores [42], was used to examine stained polystyrene nanospheres in solution. This instrument comprised an Olympus IX71 inverted microscope equipped with a 60x 1.2NA water immersion lens and an Andor iXon 888 emCCD camera (1024x1024 pixels, 13 x 13 µm). Movies were captured at a frame rate of 8.3 Hz and saved as DAX files. Excitation light from solid-state lasers (488 nm or 532 nm) was introduced to the sample via a prism coupled to a quard microscope slide, where it was totally internally reflected, exciting the aqueous sample to a depth of around 200 nm through the associated evanescent field. TIRF reduces the signal from out-of-focus fluorophores in the bulk solution and provides a low enough background to detect single fluorophores that have a high quantum yield and good photostability. The existing filters and dichroic mirrors in the emission path were optimized for the dual detection of Cy3 (550 to 620 nm) and Cy5 (640 nm long pass) emissions which are projected side-by-side on the emCCD chip. These regions span the wavelengths of interest for detecting Nile Red and DCVJ fluorescence when bound to polystyrene nanobeads.

### Raman spectroscopy

Raman spectra were collected with a Horiba Labram HR Evolution spectrometer using a 532 nm diode excitation laser at 50 mW input power. The spot size at the sample was about 2 µm. Spectra were recorded from 100 to 3800 cm^-1^, with 15 s acquisition times and 3 accumulations for each sample. A diffraction grating with a grating density of 1200 lines/mm was used for all measurements with a spectral resolution of ∼1 cm^−1^. Spectra were exported using Spectragryph [43] and analyzed using OpenSpecy using the default pre-processing and the Raman reference library of derivative-transformed spectra [44].

## Results

### Effects of solvent on fluorescence emission

The fluorescence emission of Nile Red shows a strong dependence on solvent polarity both in terms of quantum yield and Stokes shift [45,46]. DCVJ fluorescence emission also is dependent on solvent polarity but is much more sensitive to solvent viscosity [47,48]. Table 1 summarizes the wavelength maxima and relative peak intensity in several solvents measured at an excitation wavelength of 450 nm. This wavelength is near the peak for DCVJ absorption, but it is blue-shifted from the maximum absorption by Nile Red and was chosen as a convenient wavelength for comparing stained plastic samples. While both Nile Red and DCVJ show weak emission in H_2_O, Nile Red generally shows more than one order of magnitude higher emission intensity compared with DCVJ in non-polar solvents. Given that the absorbance of the DCVJ solution in isopropyl alcohol (IPA) at 450 nm was an order of magnitude higher than that of the Nile Red solution, these data indicate that the quantum yield of Nile Red is about 2 orders of magnitude greater than that of DCVJ in the solvents examined. This is in line with literature reporting the quantum yields for Nile Red dissolved in short chain alcohols are in the range of 0.4 to 0.64, while the DCVJ values are 0.002 to 0.003 [40,47–49]. In general, Nile Red showed a larger shift in emission wavelength with polarity than DCVJ, with the exception of the observed value for DCVJ in hexane which differed from a value of around 480 nm reported for other hydrophobic solvents [40]. It is possible that DCVJ dimerizes in hexane at the concentrations examined (8 µM). In a mixed solvent of hexane and isopropyl alcohol, the DCVJ emission was further red-shifted to 608 nm (Fig S5), comparable to that observed when bound to polystyrene nanospheres in water [33,34]. This observation suggests that these solvents may form a dynamic cage around a DCVJ dimer to satisfy both hydrophilic and hydrophobic interactions with the dye.

**Table 1.**
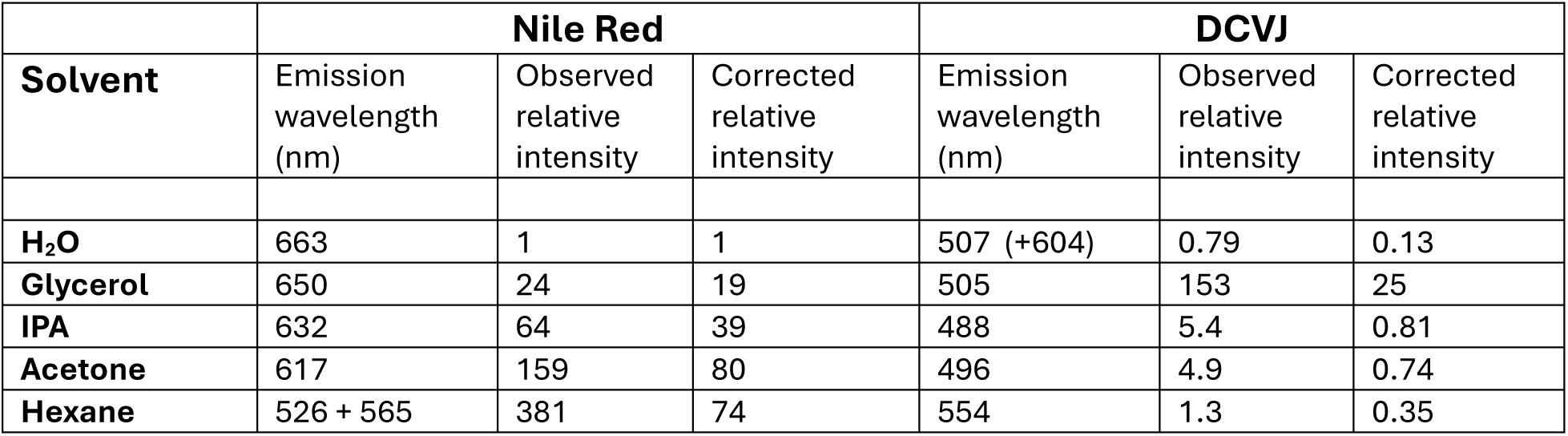
Fluorescence emission of Nile Red and DCVJ in different solvents.

| **Solvent** | **Nile Red** |  |  | **DCVJ** |  |  |
| --- | --- | --- | --- | --- | --- | --- |
|  | Emission wavelength (nm) | Observed relative intensity | Corrected relative intensity | Emission wavelength (nm) | Observed relative intensity | Corrected relative intensity |
| **H<sub>2</sub>O** | 663 | 1 | 1 | 507 (+604) | 0.79 | 0.13 |
| **Glycerol** | 650 | 24 | 19 | 505 | 153 | 25 |
| **IPA** | 632 | 64 | 39 | 488 | 5.4 | 0.81 |
| **Acetone** | 617 | 159 | 80 | 496 | 4.9 | 0.74 |
| **Hexane** | 526 + 565 | 381 | 74 | 554 | 1.3 | 0.35 |

To cover the large range in signal amplitudes, each solvent was investigated using at least two different photomultiplier voltages. The recorded intensities were then normalized in overlapping pairs to cover the complete range. Note the observed emission peak wavelength for DCVJ in hexane is an outlier, given its polarity and viscosity, possibly due to limited solubility and dimer formation. A red-shifted peak (604 nm) is also seen with DCVJ in H_2_O at high concentrations (Fig 2b). The high observed emission intensity for Nile Red in hexane is partly due to the shift in the absorption maximum towards 450 nm and also emission in a region where the detector is more sensitive. The doublet peak has been attributed to different vibronic states [50,51].

**Fig 2.**
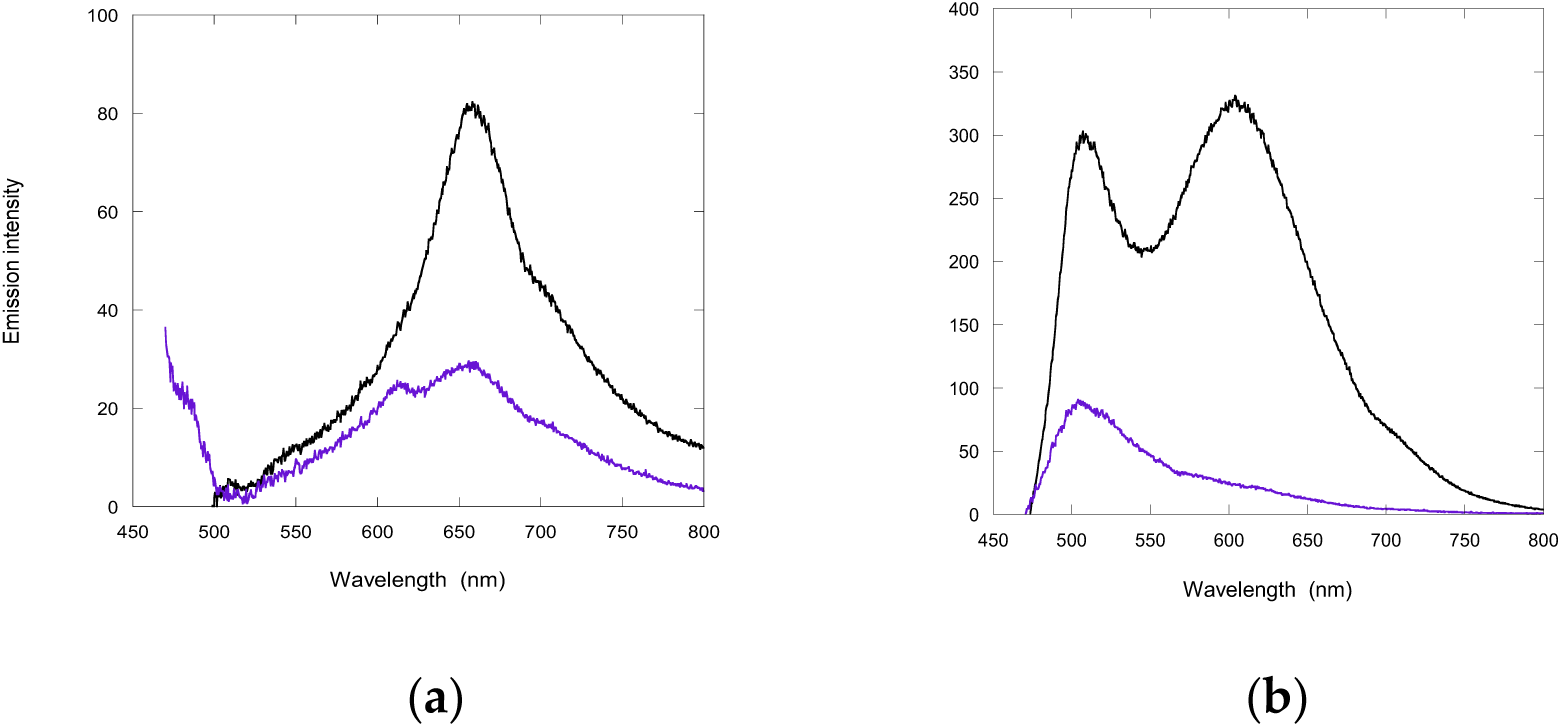
Concentration dependence of fluorescence spectrum. (a) Nile Red and (b) DCVJ fluorescence emission in H_2_O measured in a 2 mm pathlength cuvette. Blue curves = 8 µM and black curves = 40 µM. Note the ∼3-fold increase in emission for Nile Red for a 5-fold increase in concentration, indicating that dimer/aggregate formation is associated with a quench in fluorescence. In contrast, there is >10 fold increase in DCVJ emission at 604 nm for a similar concentration increase, arising from dimer formation.

Both Nile Red and DCVJ have limited solubility in H_2_O, however their behavior differs when approaching saturated concentrations. Nile Red forms aggregates that have reduced fluorescence emission at 650 nm [52], while DCVJ shows the formation of a new peak around 600 nm, likely associated with the formation of dimers [53,54]. We confirmed this behavior by comparing the spectra from 8 and 40 µM Nile Red and DCVJ in water, using a 2 mm pathlength cuvette to minimize inner filter effects (Fig 2). The emission from Nile Red solutions in H_2_O showed a decline in amplitude over a period of an hour due to aggregation.

### Staining of standard microplastic pellets

Pellets from the standard Polymer Kit 1.0 were stained with Nile Red and DCVJ under identical conditions and photographed through a 515 nm long pass filter with excitation from a 450 nm LED flashlight (Fig 3). Control unstained samples were included to test for any intrinsic fluorophores present that respond to 450 nm excitation. Both samples of PET showed strong intrinsic fluorescence, while weaker fluorescence was seen in the PEST and PA samples. The intrinsic fluorescence of these samples was more clearly seen when a 365 nm LED flashlight was used to illuminate the samples (Fig S1b). Fig 3 demonstrates the ability of Nile Red to distinguish different classes of polymer based on the color of their fluorescence emission, as reported previously [25,27,55]. In contrast, DCVJ showed very weak staining of hydrophobic polymers, such as PE and PP. Those polymers that were stained by DCVJ appeared predominantly green, particularly PA and PVC.

**Fig 3.**
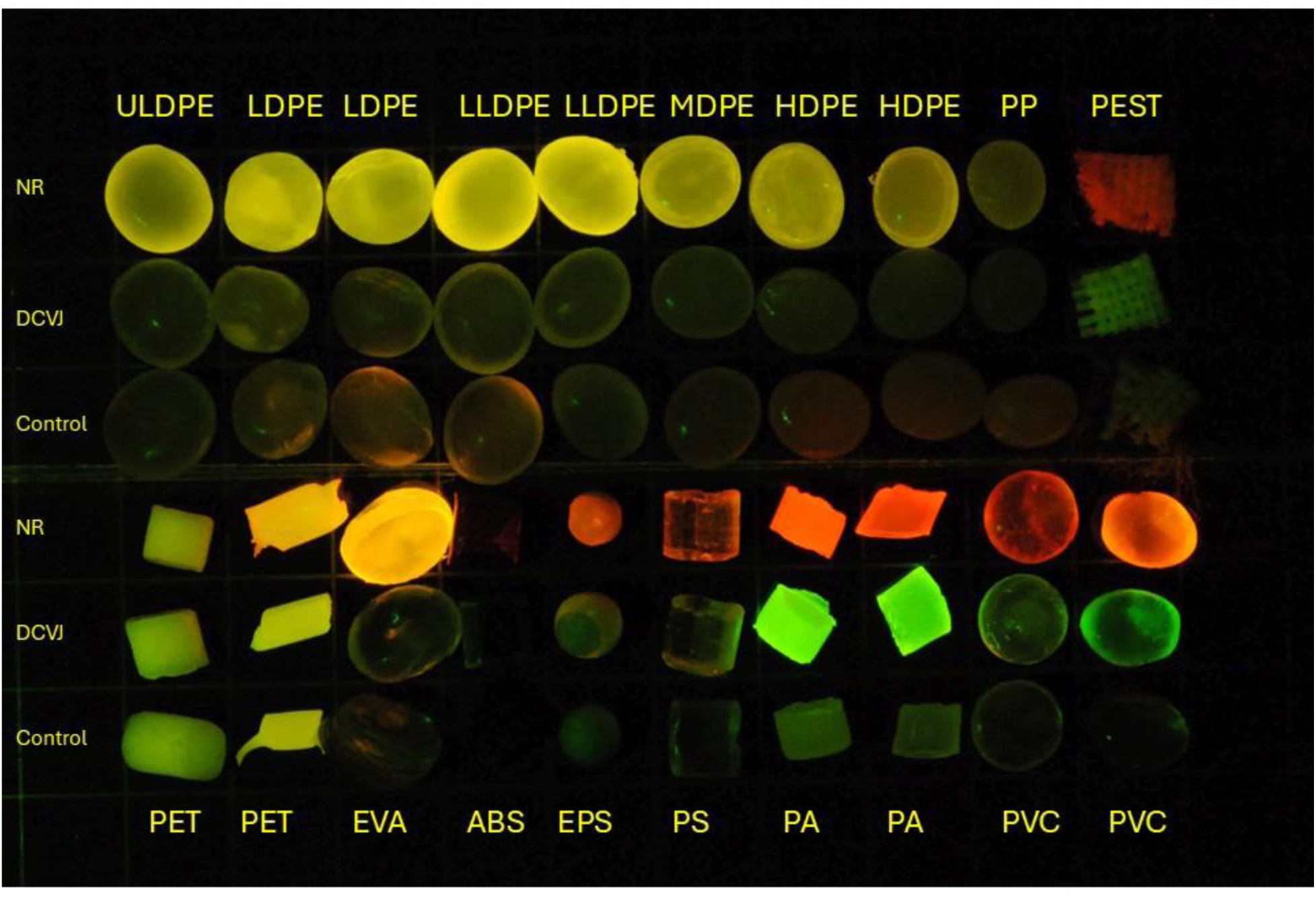
Fluorescence of stained Polymer kit 1.0 samples. Pellets stained with Nile Red (NR: 1^st^ and 4^th^ rows), DCVJ (2^nd^ and 5^th^ rows) and control unstained pellets (3^rd^ and 6^th^ rows). Excitation = 450 nm LED, emission filter = LP515 nm. Note that some of the weakly fluorescent pellets next to strongly fluorescent pellets show reflections from the latter and these regions were avoided in the RGB analysis of Table 3. See Materials section for abbreviations of plastic types.

**Table 2.** A comparison of fluorescence emission by RGB and spectral analysis.

|  |  | Nile Red |  |  | DCVJ |  |  |
| --- | --- | --- | --- | --- | --- | --- | --- |
| # | Plastic | Red chromaticity 2R/(2R+G) | Relative Intensity (2R+G) | $\lambda_{max}$ (nm) | Red chromaticity 2R/(2R+G) | Relative intensity (2R+G) | $\lambda_{max}$ (nm) |
| 1 | ULDPE | 0.47 | 27.5 | 534 + 574 |  | weak |  |
| 2 | LDPE | 0.48 | 49.2 | 534 + 575 |  | weak |  |
| 3 | LDPE | 0.47 | 45.6 | 535 + 576 |  | weak |  |
| 4 | LLDPE | 0.49 | 65.7 | 534 + 575 |  | weak |  |
| 5 | LLDPE | 0.49 | 60.4 | 535 + 576 |  | weak |  |
| 6 | MDPE | 0.48 | 33.1 | 534 + 576 |  | weak |  |
| 7 | HDPE | 0.49 | 30.6 | 535 + 576 |  | weak |  |
| 8 | HDPE | 0.49 | 19.1 | 535 + 576 |  | weak |  |
| 9 | PP | 0.47 | 8.0 | 530 + 570 |  | weak |  |
| 10 | PEST | 0.77 | 7.6 | 610 | 0.37 | 6.2 | 485 |
| 11 | PET | 0.45 | 25.6 | 611 | 0.41 | 24.4 | 485 |
| 12 | PET | 0.58 | 69.7 | 611* | 0.44 | 52.1 | 494* |
| 13 | EVA | 0.62 | 100 | broad | 0.44 | 3.9 | broad |
| 14 | ABS | 0.76 | 1.8 | 611 |  | weak |  |
| 15 | EPS | 0.70 | 15.7 | 590 | 0.47 | 5.4 | broad |
| 16 | PS | 0.68 | 10.2 | 591 | 0.41 | 7.9 | broad |
| 17 | **PA** | 0.83 | 61.6 | 615 | 0.25 | 71.6 | 497 |
| 18 | **PA** | 0.83 | 42.8 | 616 | 0.24 | 57.5 | 497 |
| 19 | **PVC** | 0.79 | 11.5 | 612 | 0.35 | 6.9 | 499 |
| 20 | **PVC** | 0.76 | 44.0 | 592 | 0.25 | 11.1 | 495 |

**Table 3.** RGB and Raman analysis of Hawai’ian environmental microplastics.

| HMP # | NR peak (nm) | NR Chromaticity (2R/(2R+G)) | Raman analysis | OpenSpecy match score |
| --- | --- | --- | --- | --- |
| 1 | 600 | 0.80 | PE | 0.85 |
| 2 | 587 | 0.74 | PE | 0.98 |
| 3 | 606 | 0.81 | PP | 0.92 |
| 4 | 598 | 0.75 | PE | 0.99 |
| 5 | 599 | 0.69 | PP | 0.93 |
| 6 | 576 | 0.69 | PE | 0.98 |
| 7 | 612 | 0.74 | PE | 0.93 |
| 8 | 531 + 572 | 0.59 | PP | 0.86 |
| HDPE | 535 + 576 | 0.52 | PE | 0.99 |

The photograph in Fig 3 was investigated further by RGB analysis using ImageJ and summarized in Table 2. The blue component was very low in all samples due to the filtering by the 515 nm long-pass filter (Fig S2). The color of the pellets was therefore quantified using the modified chromaticity ratio R/(R+G). In the case of RAW NEF images, the G component was around two-fold higher than those from JPG images, a reflection of the 1:2:1 proportion of RGGB pixel arrangement in the Bayer matrix of the camera sensor. For this reason, we report the ratio as 2R/(2R+G) when analyzing RAW images, which gives a value of 0.5 for samples that appear yellow by eye. In general, analysis of JPG and RAW images showed the same trend with different plastic samples, but JPG images were prone to saturation (i.e. pixel value = 255) of the R or G channel which distorted the chromaticity ratio. In addition, at low intensities, analysis of JPG images showed a significant value for the B channel, but this remained minor for fluorescent samples.

### Fluorescence spectroscopy of standard microplastic pellets

Spectra were recorded directly from the Polymer Kit 1.0 samples by mounting the dry pellets in the excitation beam of a Varian Eclipse spectrofluorimeter (Fig 4). Given that the pellets were slightly different shapes and had different surface textures, the extent of reflected/scattered excitation light which reached the detection port varied between samples and measurements. For strongly fluorescent samples, the contribution of stray/scattered light above 500 nm was minor, as evident from the signals from the corresponding control unstained samples. Fluorescence emission peaks were resolved for Nile Red stained samples between 534 nm (PE) and 615 nm (PA and PVC), in line with the colors seen by eye (Fig 3 and S1). In contrast, fluorescence emission from DCVJ stained pellets, where visible, was generally in the region of 500 nm and not always clearly resolved from the background light scatter. Exceptions were those from PA and PVC which showed very intense peaks at 497 nm and 495 nm respectively. Note that the photograph in Fig 3 was obtained using a 515 nm long pass filter that reduces the observed green fluorescence intensity from PA and PCV by about 4-fold. The acrylon-butadiene-styrene (ABS) sample, which appeared jet black by eye, showed very little fluorescence (note scale of graph), although there was a weak red peak with Nile Red staining. Spectra from the unstained pellets were generally dominated by scattering in the region of 470 to 500 nm, except for PET that showed strong intrinsic fluorescence extending beyond 600 nm, as noted by Konde et al [56].

**Fig 4.**
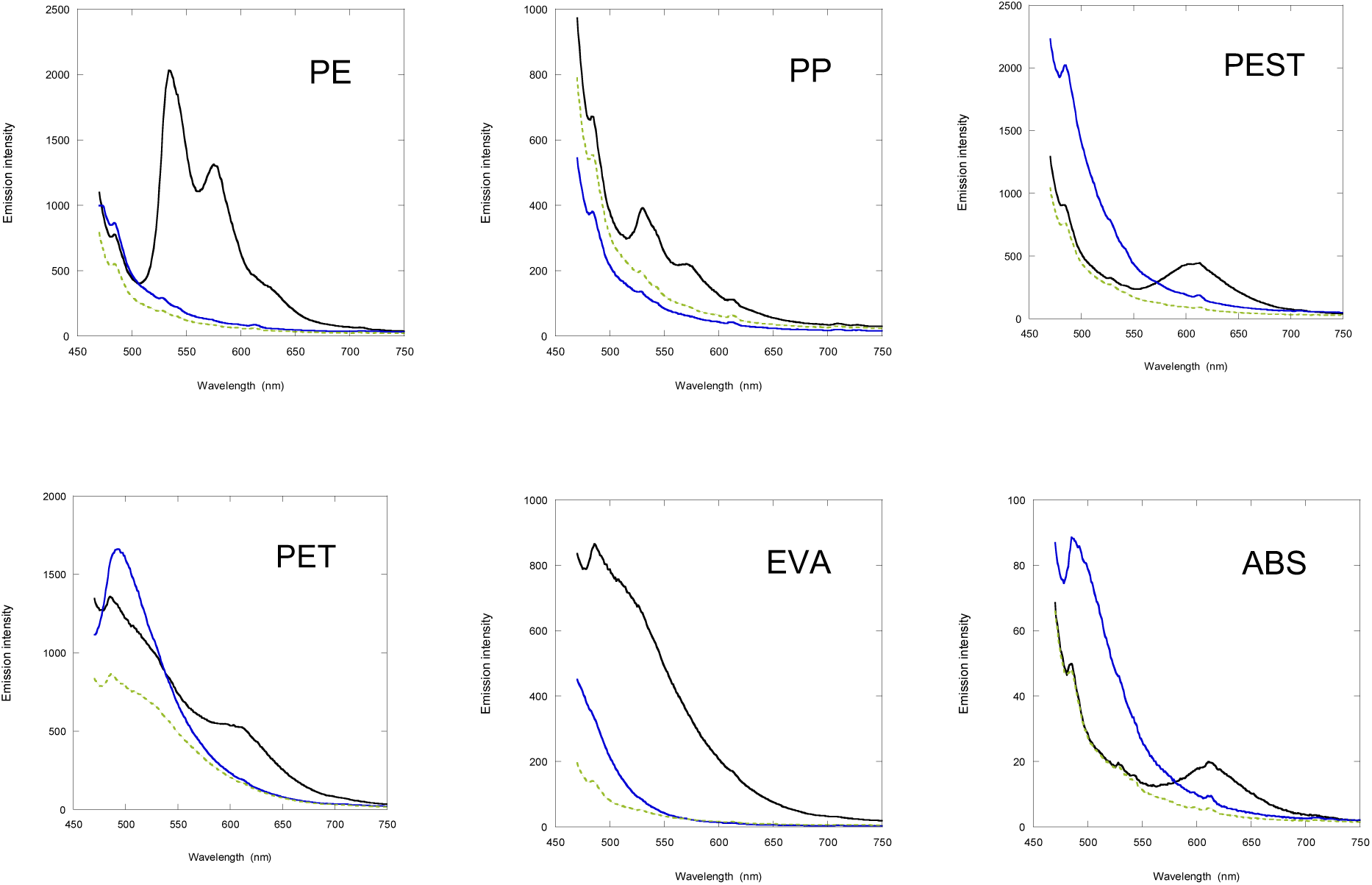
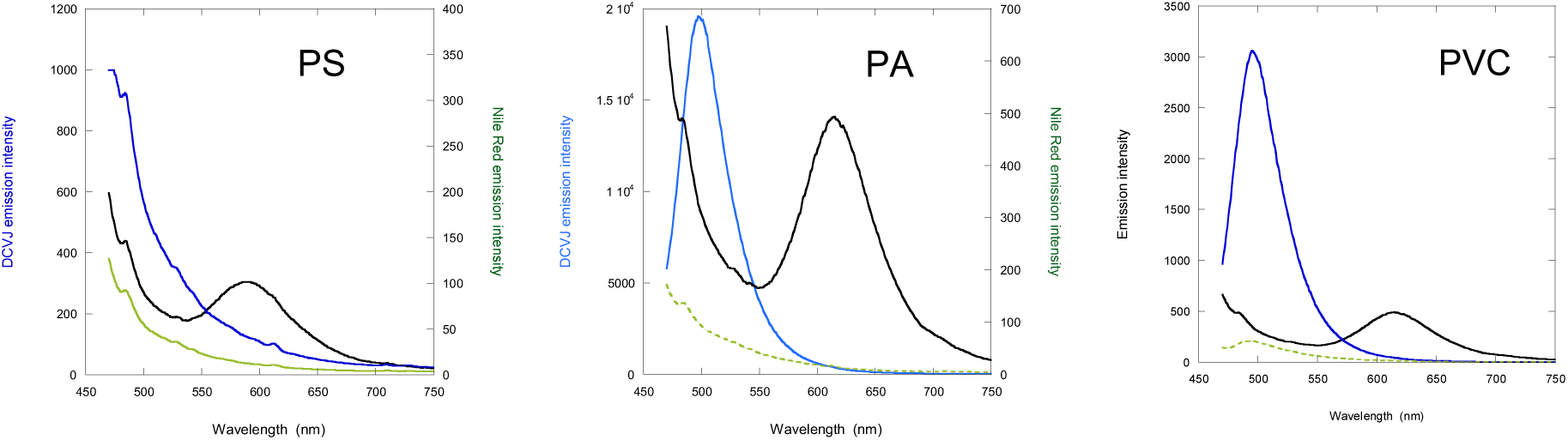
Fluorescence emission spectra for Polymer Kit 1.0 samples. Samples were stained with Nile Red- (black line) or DCVJ- (blue line), along with unstained samples (green line) to assess the scattered light and intrinsic fluorescence contribution. Spectra were measured at different photomultiplier voltages (550 to 700 V) and normalized for that corresponding to 700 V to give the ordinate values. PE spectra are only shown for the #4 LLDPE sample, as all PE spectra had a similar shape. See Materials section for abbreviations of plastic types.

The spectra from the Nile Red-stained samples are in general agreement with previous studies [55,56] and correlate with the RGB analysis of Table 2. All polyethylene pellets (ULDPE, LDPE, LLDPE, MDPE, HDPE) showed similar spectral shapes with peaks at 534 nm and 575 nm and a shoulder around 630 nm, although the overall intensities varied, likely due to the different degree of dye penetration. The double emission peaks observed with PE pellets, and weaker ones with PP, are similar to the double peaks observed for Nile Red dissolved in hexane (Table 1), where they have been attributed to vibronic states [50,51].

While the unstained control samples revealed any intrinsic fluorescence of the polymer, the scattering contribution depended on surface reflections from the pellet and its orientation in the excitation beam. Consequently, the signals could not be used for accurate baseline correction of the spectra of the stained samples, although they do provide a semi-quantitative measure of the scattering contribution. In some spectra they revealed small “ghost” peaks (55) at 528, 541 and 612 nm arising from stray light reflections from the monochromator. These peaks were insignificant compared with the broader fluorescence peaks from the stained samples, but they can become more significant at lower excitation wavelengths where the relative scattering contributions are higher (see Discussion). The ghost peaks could be removed by inclusion of a 450 nm bandpass filter in the excitation path and a 490 nm long pass filter in the emission path.

### Characterization of environmental microplastics

The same staining and analysis procedures were used to investigate microplastics collected from the environment. Plastics fragments of the order of 1 x 1 cm in size, collected from Waimānalo beach, O’ahu, Hawaii (HMP), were cut into samples of approximately 3 x 3 mm, to compare staining with Nile Red and DCVJ, together with an unstained control (Fig 5). These samples were either white or translucent, except HMP6 which was pale blue and HMP8 which was a green thread. All HMP samples showed melting with the hot needle test using a soldering iron tip at 350° C [58]. In addition to these environmental samples, fragments cut from a polystyrene Petri dish, a white HDPE food container, nylon fishing line and a cigarette filter were examined, along with a selection of microplastics from the Polymer kit as standards.

**Fig 5.**
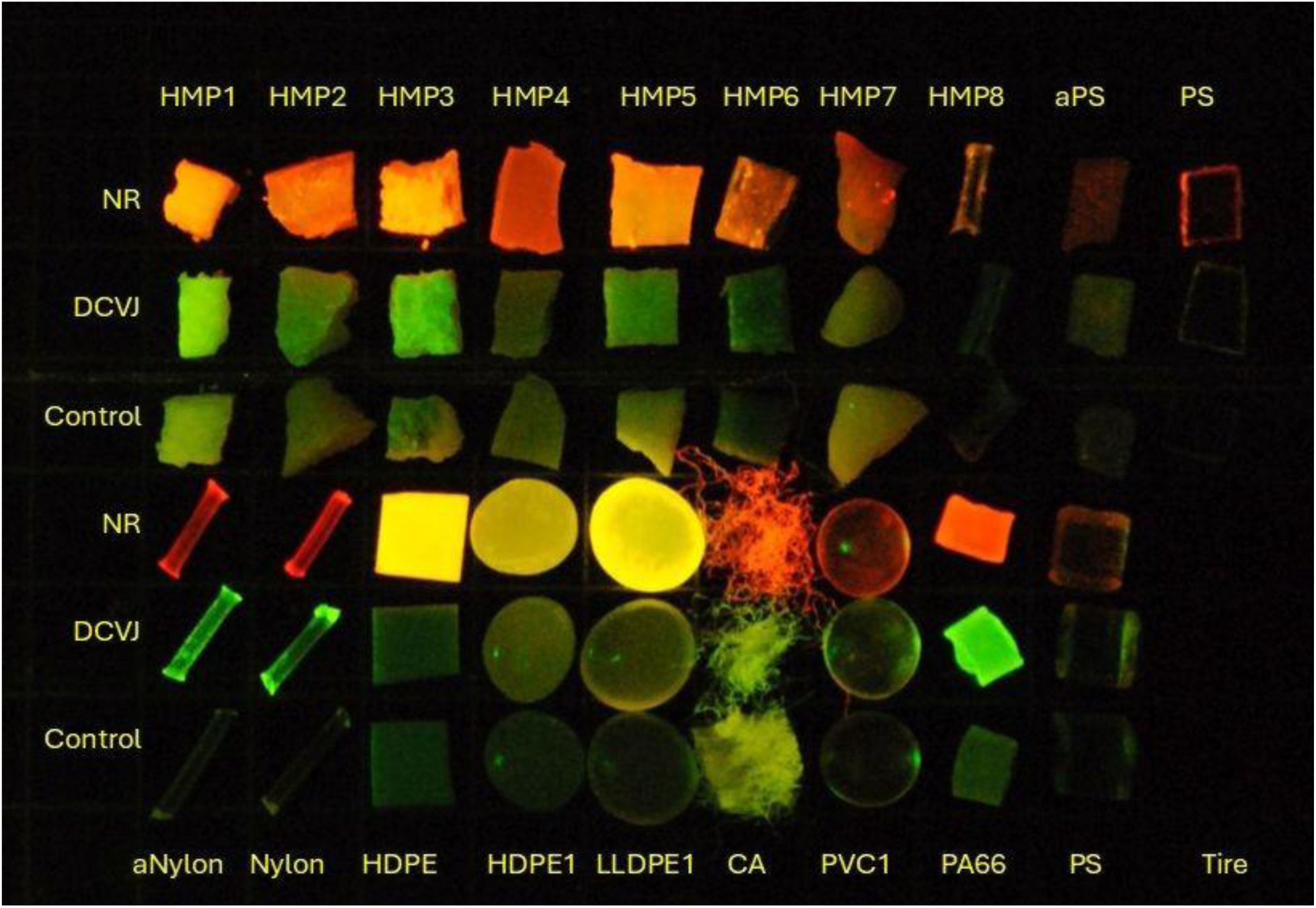
Fluorescence from environmental plastics. Staining of plastic fragments obtain from Waimānalo beach, Hawaii (HMP1 to HMP8), fragments cut from a Polystyrene Petri dish (aPS and PS), nylon fishing line (aNylon and Nylon), pristine white HDPE from container lid, HDPE1 and LLDPE1 from Polymer Kit 1.0, cellulose acetate (CA) from used cigarette filter, PVC1, PA66, PS and black tire fragment from Polymer Kit 1.0. Samples aPS and aNylon were pretreated with acetone to roughen the surface before staining. Samples stained with Nile Red (NR: 1^st^ and 4^th^ rows), DCVJ (2^nd^ and 5^th^ rows) and control unstained pellets (3^rd^ and 6^th^ rows).

Some of the unstained HMP beach samples showed relatively strong intrinsic fluorescence (e.g. HMP1 and HMP7). When stained with Nile Red, the HMP samples gave strong fluorescence signals and some had a distinct orange tinge (e.g. HMP2, HMP4, HMP6, HMP7) compared with the yellow color of standard polyethylene samples. This was confirmed by RGB analysis and emission peaks in the region of 590 to 610 nm (Table 3). HMP8 was one exception, which was a green thread-like fragment and showed characteristic double emission peaks at 531 and 572 nm indicative of a hydrophobic plastic, such as polyethylene or polypropylene. In contrast, DCVJ staining of HMPs showed dull to strong green fluorescence that was not always well-resolved from the scattering or intrinsic fluorescence background observed with unstained samples.

While samples HMP1 to HMP7 were initially identified as hydrophilic plastics, based on their Nile Red fluorescence staining, Raman analysis showed peaks characteristic of polyethylene and polypropylene for all HMP samples (Table 3 and Fig 6). This suggests that environmental factors, such as oxidation, and/or additives were affecting the interaction with the dye. Examination of the Raman spectra showed only very minor peaks in the C=O region, suggesting oxidation was not extensive [59]. Freshly cut surfaces of the HMP samples also stained orange indicating the cause was not restricted to the exposed surface layers. The most likely explanation is the interaction of Nile Red with hydrophilic additives used in the production of the plastics [60]. Some samples (e.g. HMP1 and HMP3) also showed strong green fluorescence with DCVJ staining likely due to the same hydrophilic additives. The relatively pristine white HDPE sample from a container lid stained a yellow color with Nile Red (2R/(2R+G) = 0.60 and showed emission peaks at 535 and 575 nm, as expected.

**Fig 6.**
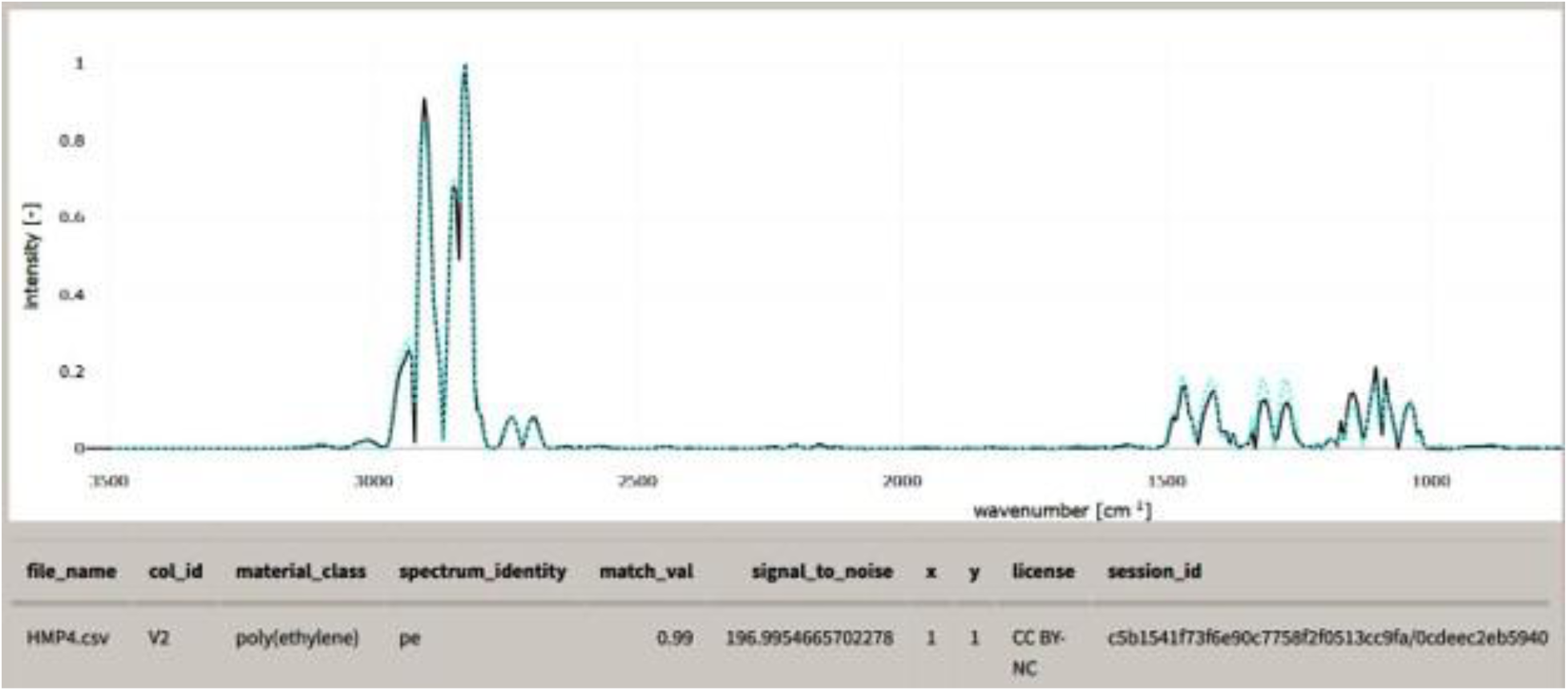
Raman spectrum of HMP4 and analysis using OpenSpecy. [44].

The staining of polystyrene is of particular interest, given the ability to detect PS nanoplastics using DCVJ [33,34]. One set of the test fragments (aPS) cut from a pristine polystyrene Petri dish were pre-treated with acetone for 1 minute to roughen the surface and to compare with untreated transparent polystyrene. Both were stained with Nile Red and DCVJ in 25% acetone, 75% water solution [27] but the untreated samples remained transparent. After staining, it is evident that the acetone-pretreated samples stained throughout the exposed surfaces whereas the untreated samples only stained at the freshly cut surfaces (Fig 5, samples aPS cf. PS). The DCVJ stained polystyrene was dominated by green fluorescence, in contrast to the 620 nm emission reported for polystyrene nanospheres [33,34]. In a separate experiment, polystyrene Petri dishes were stained in the presence of 98% acetone, rather than just pretreatment, which gave more intense fluorescence emission (Fig S7). Analysis in the spectrofluorimeter showed an emission peak for Nile Red staining at 583 nm (Fig Sd). In contrast, DCVJ staining gave a peak at 486 nm after correction for background scattering, with no indication of any red emission. The nylon fishing line samples also stained more strongly at the cut ends but in this case acetone pre-treatment made little difference to the overall staining (Fig 5, samples aNylon cf. Nylon).

Cellulose acetate fibers from a used cigarette filter showed strong red fluorescence with Nile Red staining (Fig 5, CA), as we reported previously [29], and strong green fluorescence with DCVJ. The black tire fragments from the Polymer Kit (Fig 5, Tire) failed to yield any signal.

### Characterization of polystyrene nanoplastics

In view of the previous studies showing that polystyrene nanobeads stained with DCVJ give a distinct fluorescence emission at 620 nm [33,34] and the above results with millimeter-sized polystyrene samples that showed no sign of red emission, we investigated the staining of 70 nm polystyrene beads (PS70). Addition of PS70 beads to Nile Red in water showed a quench in fluorescence and a blue shift from around 657 to 620 nm (Fig 7a). In contrast, 8 uM DCVJ in water showed weak emission in the red due to dimer formation [54], which became dominant and peaked at 620 nm on addition of 70 nm PS beads (Fig 7b), in line with previous studies [33,34]. Moraz & Breider [34] noted that the emission at 620 nm was dependent on the diameter of the beads and the signal showed little increase on increasing the bead concentration with 300 nm diameter PS beads compared with ≤100 nm beads. This property could reflect the length of individual polymer chains compared with the dimension of the bead, which would affect the possible packing arrangements of the chains.

**Fig 7.**
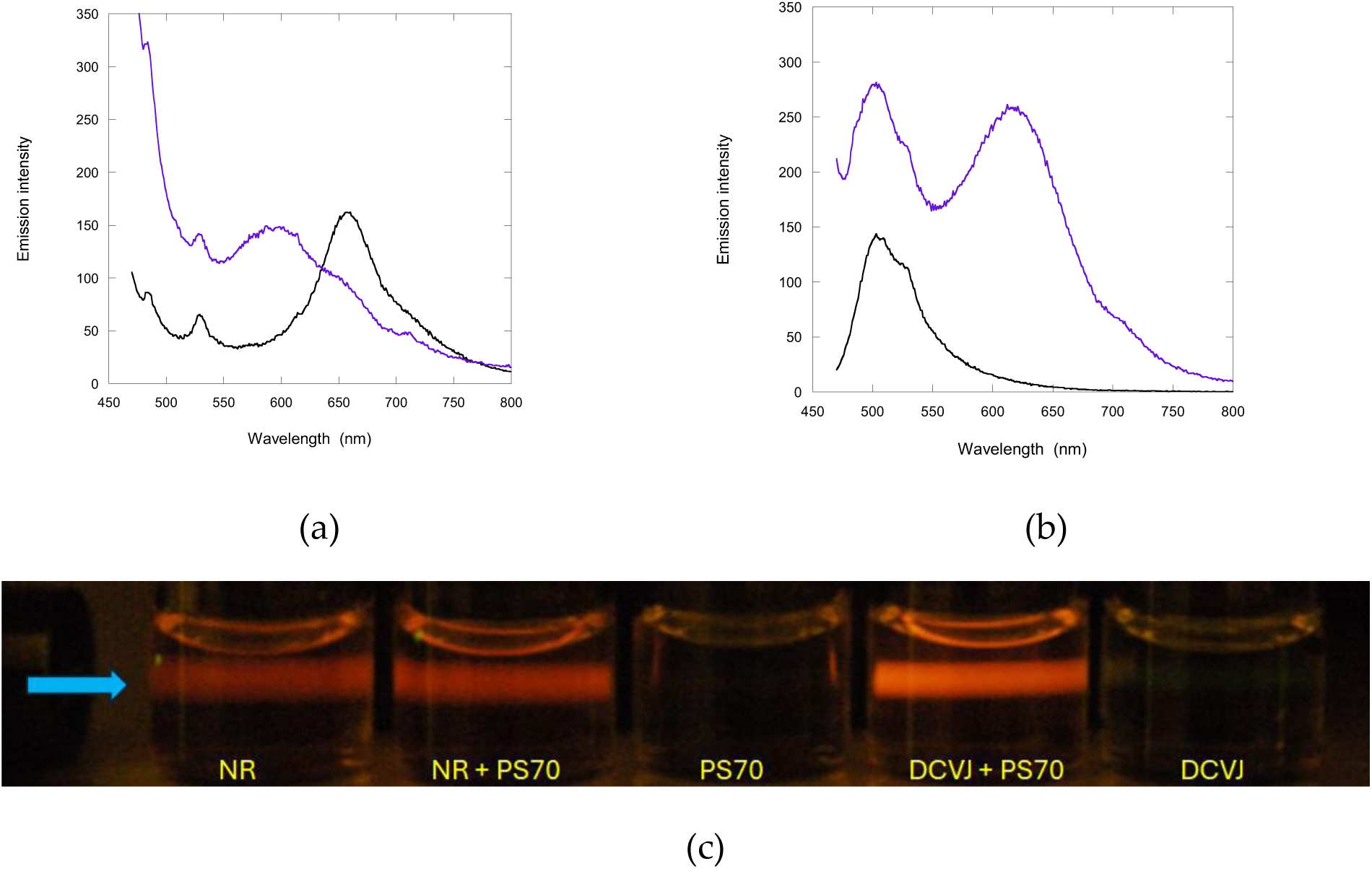
Fluorescence emission from stained 70 nm polystyrene beads (PS70). (a) Emission spectra of 8 µM Nile Red (NR) in H_2_O (black) and on addition of 10 µg/mL PS70 beads (blue). (b) Emission spectra of 8 µM DCVJ in H_2_O (black) and on addition of 10 µg/mL PS70 beads (blue). Note the increase in signal below 550 nm on PS70 addition, in both (a) and (b), due to breakthrough of stray scattered light. (c) Photograph of a 450 nm laser beam passing through the same samples in aligned 1 cm pathlength cells, plus a control for PS70 alone showing the absence of intrinsic fluorescence, taken with a Nikon D3000 dSLR and Zeiss 515nm long pass filter. Analysis of RAW images gave 2R/(2R+G) chromaticity values of NR = 0.78, NR+PS70 = 0.77, DCVJ+PS70 = 0.74 and DCVJ = 0.27.

The same characteristics of Nile Red and DCVJ staining of 70 nm polystyrene beads were revealed by photography, when a 450 nm laser beam was shone through the sample cuvette (Fig 7c). Note that the camera images were obtained using a 515 nm long pass filter that suppresses the green signal from free DCVJ. Also, the camera has a built-in infrared filter (Fig S2a) that suppresses the red (>650 nm) fluorescence from the free Nile Red. The images of Fig 7c are consistent with the spectral analysis in showing that, at the concentrations used, the DCVJ-stained PS70 beads gave a slightly higher orange-red intensity than the Nile Red-stained beads, while DCVJ alone emitted in the green region.

Samples with the same composition as in Fig 7 were then examined directly by fluorescence microscopy. Initial measurements were made with two microscopes: (i) a custom-built microscope with a long working-distance 20X Mitutoyo objective lens, assembled with the longer-term goal of incorporating stimulated IR Raman optics (Fig 8) and (ii) a consumer-grade Amscope 120 microscope with a Nikon D3000 dSLR and 515nm long pass emission filter (Fig S8). Observations were first made on dried samples to minimize sample thickness and reduce diffusion during image acquisition. When 5 µL aliquots of samples containing 70 nm PS beads were evaporated on a slide, the beads tended to aggregate in a ring around the edge of the spot, while the center contained individual particles (Fig 8 and S8). Dried samples containing Nile Red or DCVJ alone, produced aggregates that were more evenly distributed across the area and were generally larger than most of the particles observed in the presence of PS70. Many of the individual particles observed with samples containing stained 70 nm polystyrene beads had a diameter consistent with the calculated diffraction limit for the objective lens (i.e. with a NA = 0.4 objective lens and 600 nm light, the limit (= 1.2 * λ/2NA) is 900 nm), suggesting they represented single beads or aggregates of just a few beads (Fig 8b).

**Fig 8.**
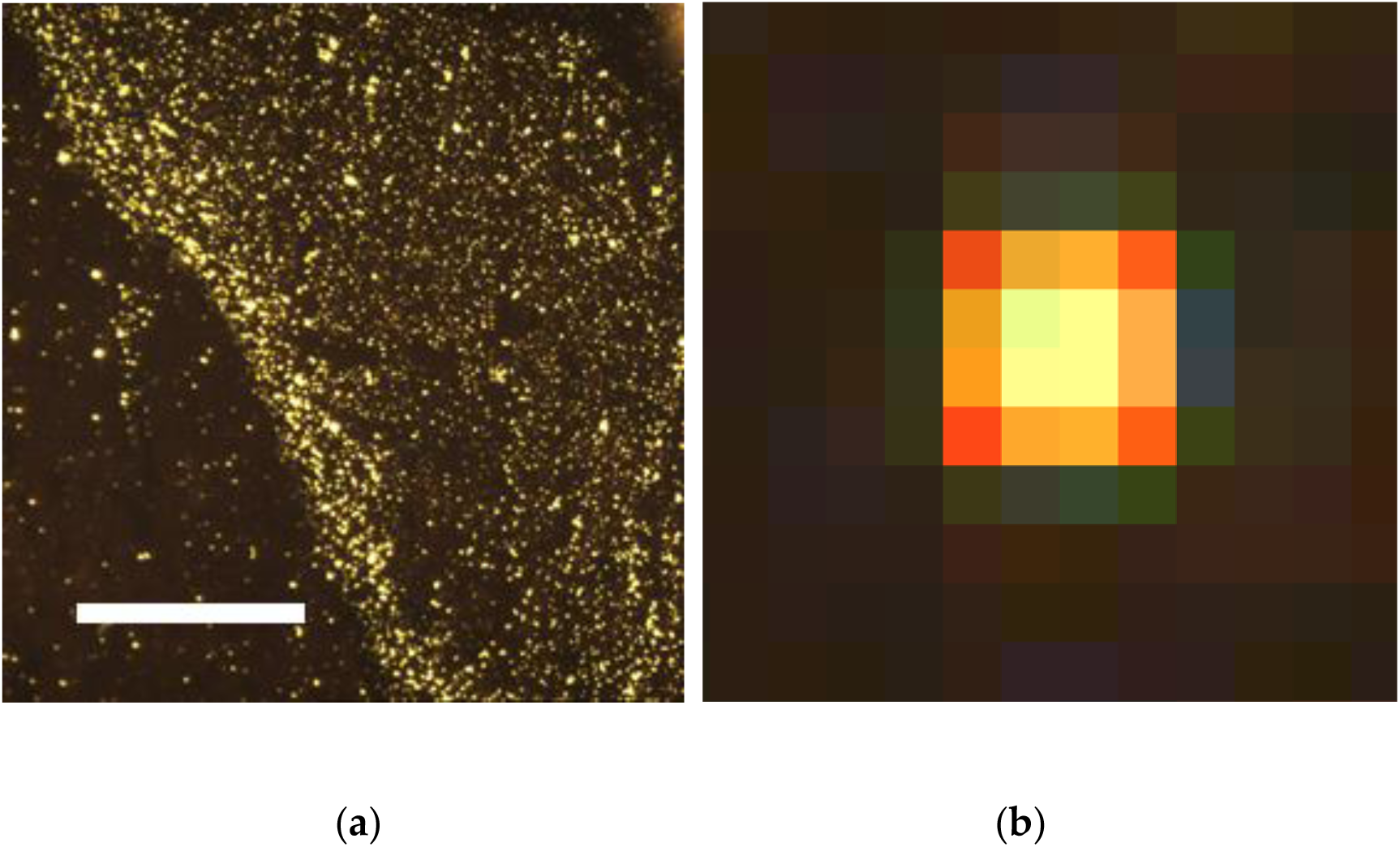
Microscopy of Nile Red-stained polystyrene nanobeads. (a) Fluorescence image of the edge of a dried sample of Nile Red-stained 70 nm polystyrene beads taken with a 20x 0.42 NA Mitutoyo objective lens and FLIR Blackfly CMOS camera (0.24 s exposure). Excitation = 450 nm, emission filter = OG530 (Figure S2a). Scale bar = 100 µm. (b) Enlarged image of a single particle selected from the inside of the evaporated boundary. The spot size is diffraction-limited by the objective lens while the particle itself is located near the center of the image, giving a near symmetrical pixel intensity distribution. Pixel image size = 0.3 x 0.3 µm.

The detection system of the Amscope/Nikon dSLR combination (Fig S8) allowed direct comparison with the images obtained for the same samples while in solution (Fig 7c). The chromaticity ratio, 2R/(2R+G), for Nile Red-stained PS beads, changed from 0.77 to 0.5 and DCVJ-stained PS beads from 0.74 to 0.58 on drying. The emission peak at 620 nm from DCVJ-stained PS70 beads in solution (Fig 7b), likely associated with excimer formation, appears critically dependent on having water molecules in the vicinity of the dyes. This characteristic was noted by Moraz and Breider [34], who found that an increase in methanol/water ratio caused the emission peak to shift from 620 nm to 570 nm. In contrast, samples containing Nile Red or DCVJ alone showed a different chromaticity trend on drying, with Nile Red remaining similar at 0.78, while DCVJ increased from 0.27 to 0.82. The latter result is expected from the data in Fig 2b and the observations of Gavvala et al. [54], showing that as the DCVJ concentration increased from 8 µM to saturation during evaporation, red-emitting dimers and aggregates would form.

To obtain images of stained PS70 polystyrene beads with an increased signal-to-background ratio, aqueous samples made with the same composition as in Fig 7 were examined using total internal reflection fluorescence microscopy. This method detects emission from particles within about 200 nm of the quard slide surface and reduces signals from out-of-focus material in the bulk solution [61]. The Nile Red-stained PS70 bead sample, when excited at 532 nm (13 mW), showed numerous spots, many of which were diffraction-limited in diameter (Fig 9a). Given the resolution limit of the 1.2 NA objective lens of ∼ 0.3 µm, which corresponds to 2 x 2 pixels in this optical set-up, these spots could correspond to single PS beads, or aggregates containing ≤ 10 beads (Fig 9g,i). However, a significant number of spots were larger than the diffraction-limited diameter likely due to aggregation of the beads and/or the dye (Fig 9h). Similar but less intense images were obtained when the samples were excited at 488 nm. Video records showed about 10% of the small spots appeared and disappeared over a 12 s period, as a result of diffusion into and out of the evanescent field. The larger aggregate spots were static on this time scale, suggesting these particles were stably adhered to the quard surface. Samples of Nile Red alone also showed many spots (Fig 9b), most of which were larger than the diffraction limit, although the total number of spots was less than in the presence of PS70.

**Fig 9.**
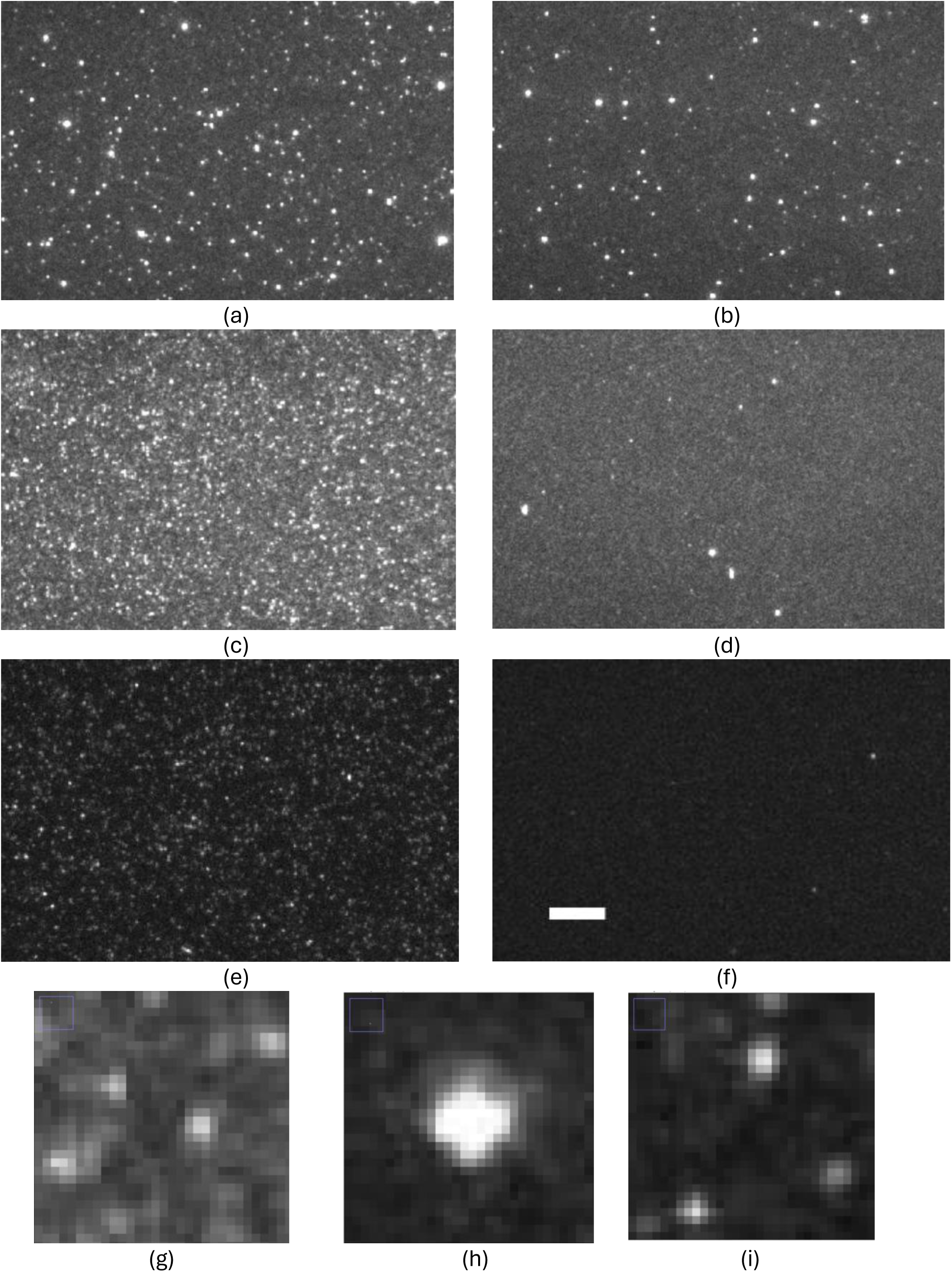
Total internal reflection fluorescence microscope images of stained 70 nm polystyrene nanobeads. Images taken with a 60x, 1.2 NA water-immersion lens. (a) 8 µM Nile Red plus 10 µg/mL PS70 in H_2_O excited at 532 nm (13 mW) with 640 nm long pass emission filter. (b) 8 µM Nile Red alone. (c) 0.8 µM Nile Red plus 10 µg/mL PS70 in 90% H_2_O, 10% isopropyl alcohol excited at 532 nm. (d) 0.8 µM Nile Red alone. (e) 8 µM DCVJ plus 10 µg/mL PS70 in H_2_O excited at 488 nm (17 mW) with 550 to 620 nm band pass emission filter. (f) 8 µM DCVJ alone. Scale bar = 10 µm in panels (a) to (f). Unstained PS70 gave no significant signal with either 532 or 488 nm excitation. Panels (g) to (i) are enlarged sections taken from panels (c), (d) and (e) to show individual particles (pixel size = 0.18 x 0.18 µm) where (g) Nile Red + PS70, (h) Nile Red alone and (i) DCVJ + PS70.

At the concentration used (8 µM), Nile Red shows extensive aggregation in water which complicates the detection of the stained nanoplastics, as previously noted by Chatterjee et al. [62]. Although the fluorescence emission per Nile Red molecule is reduced upon aggregation (Fig 2a), the spots appear bright on a darker background because of the local high concentration of molecules within the aggregate. Reducing the Nile Red concentration 10-fold, adding 10% isopropyl alcohol and minimizing the time from mixing to observation to about 5 minutes, reduced the number of spots seen with the dye in the absence of PS70 beads (Fig 9d) but those detected were still generally larger than the diffraction limit (Fig 9h). DCVJ-stained PS beads, when excited at 488 nm (17 mW), showed less intense emission that required a 4-fold increase in the emCCD camera gain to give a comparable signal to those stained with Nile Red (Fig 9e, i). However, the DCVJ alone control showed many fewer spots (Fig 9f), indicating much less of a problem with aggregation of the free dye. This is consistent with Fig 2b showing at 8 µM in water, DCVJ remains largely monomeric, as judged by the lack of a significant 604 nm dimer emission peak. For direct visualization of polystyrene nanobeads by fluorescence microscopy, DCVJ staining therefore offers a potential advantage over Nile Red in having fewer false positives, despite the weaker signal due to a lower quantum yield.

## Discussion

Staining of the standard plastic pellets in the Hawai’ian Polymer kit 1.0 revealed that Nile Red is a better stain than DCVJ for identification of plastic types, both in terms of its sensitivity and potential ability to distinguish different classes based on their hydrophobicity. In addition, Nile Red generally shows a larger Stokes shift than DCVJ, which reduces the background signal from stray scattered light. Previous work reported that Nile Red has a quantum yield of 0.4 in methanol, 0.75 in acetone [49,63] and is two orders of magnitude lower in water, whereas DCVJ has a much lower quantum yield of 0.0022 in methanol [40] rising to 0.1 in glycerol [47]. These findings are consistent with the data of Table 1 and demonstrate that Nile Red is primarily polarity-sensitive, while DCVJ is more viscosity dependent. Also, these results suggest that Nile Red will produce more intense signals than DCVJ when bound to microplastics. However, observed intensities are also dependent on the wavelength sensitivity of the detector employed. Both the sensor in the color camera and photomultiplier in the fluorimeter used in our studies are more sensitive to green light than red and, in part, account for the strong emission peaks observed for PA and PVC when stained with DCVJ (Fig 2). Another factor is the choice of excitation wavelength of 450 nm which corresponds to the peak for DCVJ absorbance, while it is at the blue edge for Nile Red excitation. In some environments and concentrations, DCVJ does show a large Stokes shift which appears to be associated with dimer/excimer formation [53,54]. This property is observed with DCVJ-stained polystyrene nanobeads with diameters <300 nm [34] and certain solvent mixtures which combine hydrophobic and polar media (Fig S5) and is discussed further below. Both Nile Red and DCVJ stain natural material such as lipids and fats, which accounts for their use as biological stains a decade or more before their application to microplastics [64,65]. This property requires pretreatment of microplastic samples with oxidizing conditions to remove organic matter in many environmental and most biological samples.

In addition to comparing two fluorescent dyes, we also used two methods to record the fluorescence; photography and spectroscopy. The latter gives higher resolution information, as demonstrated by the characteristic double vibronic emission peaks observed with Nile Red in apolar solvents (e.g. hexane) and bound to hydrophobic plastics (e.g. polyethylene). Photography can be carried out with low-cost equipment and is therefore accessible to community science projects focused on monitoring environmental microplastics. In general, there is reasonable agreement between the two approaches (Fig S6). However, the determination of a chromaticity value or peak emission wavelength does not always enable the identity of the plastic polymer type because of the influence of additives (Table 3). Nevertheless, fluorescence detection remains a useful research tool for prescreening samples before further analysis using FTIR or Raman spectroscopy.

The interaction between fluorescent dyes and plastics depends on a number of factors, such as incubation time, temperature and solvent. Furthermore, different forms of plastic may require different temperatures and solvents to soften the polymer to aid dye penetration. Several studies using Nile Red have recommended specific conditions in an attempt to make comparisons between different studies easier, although no standard method has been agreed upon [25,27,55,56,66,67]. To add to this complication, the methods of fluorescence detection also vary and have an equally significant impact on the result. A more practical way forward is to include standard plastics samples so that the effect of staining conditions and instrumentation can be assessed. The Polymer Kit 1.0 from the Hawai’i Pacific University [38] provides one such standard, although other sources are available [68]. Previously, Gao et al [67] used the Hawai’ian Polymer kit to test staining conditions with Nile Red and reported that all the plastic samples showed stronger signals in the red than green channels, which appears to differ from our studies, where some hydrophobic plastics (e.g. PE) show a more significant green component. This difference can be explained by differences in instrumentation, such as the light source for excitation. We used a constant excitation wavelength of 450 nm for analysis, whereas Gao et al [67] switched to green excitation (545 ±25 nm) for the red emission channel. Their light source, an X-Cite 120Q fluorescence lamp, is a mercury arc source with a strong emission line at 546 nm which would enhance the observed red fluorescence. Differences in the bandpass of the emission filters, as well as the spectral sensitivity of the detector, would also contribute to the green/red weighting. It is therefore useful to include these specifications in any report (Fig S2).

Our choice of 450 nm as the standard excitation wavelength for Nile Red and DCVJ is based on several factors, including the availability of consumer LED flashlights. This wavelength is at the blue edge of the Nile Red absorption spectrum and near the peak for that of DCVJ. Given that the molar absorption coefficients are: Nile Red, 19,600 M^−1^ cm^−1^ at 552 nm in DMSO [39] and DCVJ, 62,000 M^−1^ cm^−1^ at 454 nm in methanol [40], 450 nm favors the excitation of DCVJ. Excitation at 450 nm allows the full span of Nile Red emission in different solvents and polymers to be assessed, whereas 552 nm excitation would give a high scattering contribution in the green region and obscure emission from polymers such as PE. Other studies have used 405 nm to excite Nile Red, which corresponds to a near minimum in the absorption spectrum of Nile Red. Prasad et al. [55] reported specific narrow emission peaks at 485, 529 and 552 nm with several plastic samples, but these are likely stray scattered light (ghost) peaks from the monochromator grating [57]. We have observed such peaks using the same make of Varian fluorimeter and a blank glass slide angled at 40° to reflect some of the stray excitation light into the detector (Fig S4d). On the other hand, Konde et al. [56] used a high power 405 nm laser (no monochromator), in conjunction with emission long pass filters and high concentrations of Nile Red dye to detect emission spectra which closely resembled our spectra obtained at 450 nm excitation. Ultraviolet excitation has also been used with Nile Red [27], but these shorter wavelengths are more likely to excite intrinsic fluorescence found in many environmental microplastics [69,70], compounding the emission from the added dye (Fig S1b).

At high concentrations (>8 µM), DCVJ in water reveals the appearance of a red-shifted emission peak (Fig 2b) that has been ascribed to formation of dimers or higher aggregates [53]. Given that the fluorescence lifetime of DCVJ in water is of the order of 0.2 ns [54], such dimers must preexist because, according to the Smoluchowski diffusion equation [71], monomer collisions occur on the µs timescale at these concentrations. However, when bound to macrocyclic hosts such as ϒ-cyclodextran, nearby DCVJ molecules may form true excimers [29]. It is possible that the extensive looping of the long (> 1 µm) polystyrene polymer chains, when confined to within a 100 nm diameter or smaller bead, creates cavities that can accommodate closely-spaced DVCJ molecules, favoring the formation of excimers. On the other hand, in larger microplastics, the polystyrene polymer chains can pack side-by-side and have a fewer proportion of the loops that favor excimers, accounting for the lack of 620 nm emission. The precise wavelength of the excimer emission is strongly solvent dependent. The observed peak at 620 nm requires a contribution from water molecules and shifts to 570 nm in the presence of methanol [11]. We also observe a blue shift when DCVJ-stained 70 nm polystyrene beads, prepared in water, dry out on a microscope slide. The critical dependence of the DCVJ red emission on both the binding site geometry and solvent accessibility, indicates it is not a robust probe for nanoplastics in general.

Several studies have been directed towards the detection of sub-micron sized plastic particles using Nile Red [62,72–75] or DCVJ [33,34] staining but, as far as we know, no direct comparison has been published. At the concentrations and excitation wavelength used in Fig 7, the emission intensities of 70 nm polystyrene beads stained with Nile Red and DCVJ are comparable. However, the absorbance of the DCVJ sample at 450 nm was an order of magnitude higher than that of Nile Red solution, indicating the DCVJ quantum yield, when bound to polystyrene, is significantly lower. Nevertheless, the sensitivity of polystyrene bead detection depends not just on the quantum yield of the bound state, but also the minimal contribution from free dye to the background signal. While the TIRF microscope is capable of detecting single fluorophore molecules [42], the emCCD camera gain used here was about 10-fold less than required. In addition, monomeric dye molecules would diffuse on the acquisition time scale (0.12 s) and contribute to a homogeneous background signal. Here, DCVJ has a potential advantage in that the free dye emits predominantly at 507 nm, well separated from the bound dye emission (620 nm), whereas with Nile Red, both free and bound dye emit in the 620 to 660 nm region (Fig 7). In addition, DCVJ has less of a tendency to aggregate at the concentrations used, hence reducing the problem of false positive spots seen with Nile Red. Nevertheless, the absolute quantum yield of photoemission remains an important factor when detecting single nanoplastics, where higher excitation intensities are required to give detectable emission signals and are ultimately limited by photobleaching events. The development of high quantum yield variants of DCVJ [76] may offer a real advantage for nanoplastic detection as they are more sensitive to both solvent polarity and viscosity.

## Conclusions

The following are the key conclusions from this work:

- While standard protocols are helpful in comparing results from different studies, inclusion of standard plastics is essential to control for different spectral sensitivities of the instrumentation used.
- Nile Red is a generally better stain than DCVJ for microplastics because the emission is more intense for most kinds of plastics and shows a greater Stokes shift which reduces light scattering contributions.
- Nile Red staining shows a larger range of peak emission wavelengths (534 to 615 nm) with different microplastic types than DCVJ (484 to 499 nm), potentially allowing for the identification of the general type of plastic (hydrophobic or hydrophilic).
- Additives in plastic can shift the color and emission wavelengths, so other methods (e.g. Raman spectroscopy) are required to confirm the plastic composition.
- Total internal reflection fluorescence microscopy can detect individual, or small clusters of stained 70 nm polystyrene nanobeads in solution.
- While polystyrene nanobeads stained with Nile Red give a stronger emission than with DCVJ, the latter show a 620 nm emission peak (due to excimer formation) that is well separated from free monomeric DCVJ molecules and hence gives a lower background signal. In this regard, Nile Red suffers from the formation of small aggregates (≤ 1µm) in water which are difficult to distinguish from the stained nanobeads.

## Supporting information

S1 Supplementary Figs 1-8 (pdf)

## Acknowledgments

We thank Michael Stone for access to laboratory facilities. We are grateful to Rebecca Braslau and David Jameson for discussions.

## Author Contributions

Conceptualization: Clive R Bagshaw

Formal Analysis: Madison Wallner, Clive R Bagshaw

Investigation: Madison Wallner, Justin Diaz, Amelia B Labbe, James J. Jacob, Quentin Williams, Clive R Bagshaw

Methodology: Madison Wallner, James J Jacob, Clive R Bagshaw Project Administration: Clive R Bagshaw

Resources: Madison Wallner, Amelia B Labbe, James J Jacob, Quentin Williams, Adina Paytan

Supervision: Adina Paytan, Clive R Bagshaw Validation: Madison Wallner, Clive R Bagshaw Visualization: James J Jacob, Clive R Bagshaw

Writing – Original Draft: Madison Wallner, Clive R Bagshaw

Writing – Review & Editing: Madison Wallner, Justin Diaz, Amelia B Labbe, James

J. Jacob, Quentin Williams, Adina Paytan, Clive R Bagshaw

