## Supplementary Figures S1-S8 for "The comparative strengths and limitations of Nile Red and 9-(dicyanovinyl)-julolidine (DCVJ) fluorescent dyes for detecting microplastics and nanoplastics"

**Wallner et al. Supplementary Figures**

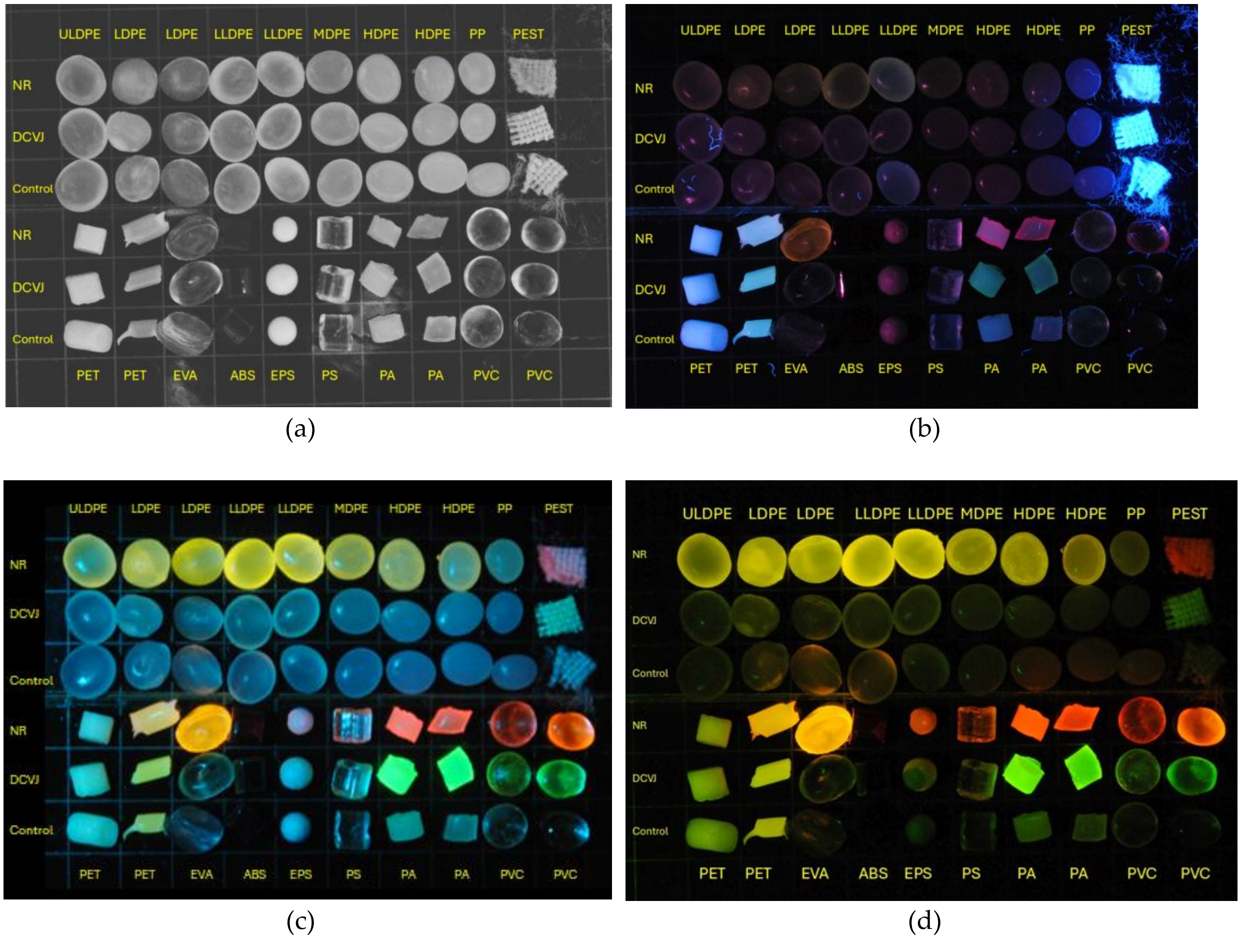


Fig S1. Polymer Kit 1.0 pellets photographed under (a) reflected white light, (b) 365 nm uv light from an LED flashlight to show the strong intrinsic fluorescence of some samples (PEST and PET) and (c) 450 nm excitation with an acrylic 490 nm long pass emission filter which transmits about 2% of the LED excitation light and gives a larger scattering contribution. (d) Copy of main text Fig 3 obtained using a 515 nm Zeiss long pass emission filter which reduced the blue scattered light to about 0.2%.

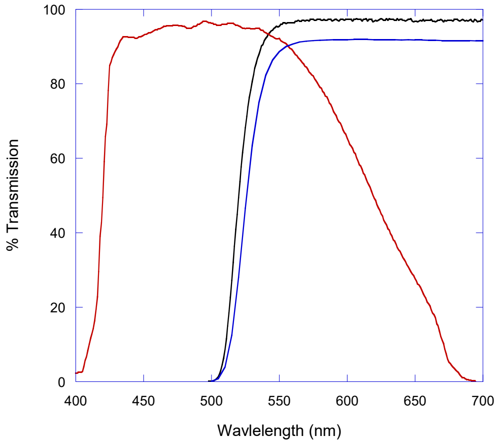


(a)

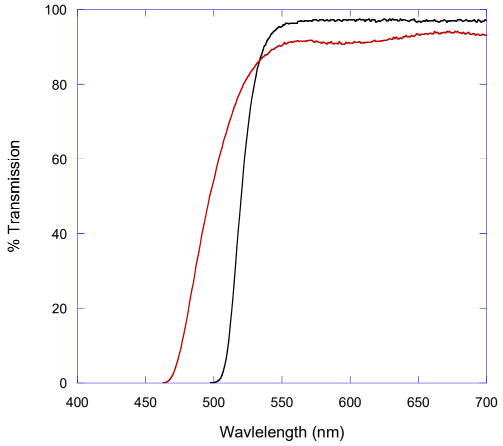


(b)

Fig S2. Transmission curves for filters used in photography. (a) Zeiss LP515 (black), Schott OG530 (blue), Built-in Nikon D3000 uv-IR filter (red). The transmission values at 450 nm are: Zeiss LP515 < 0.02% and Schott OG530 < 0.004%. However, because of the finite bandwidth of the 450 nm LED output (~ 35 nm full-width, half-maximum), the LP515 transmits about 0.2% of the excitation light. (b) Comparison of the Zeiss LP515 filter (black) with an acrylic LP490 filter (red) used in Figs S1d and S1c respectively.

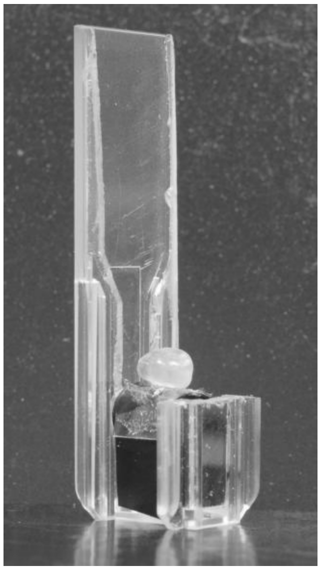

(a)

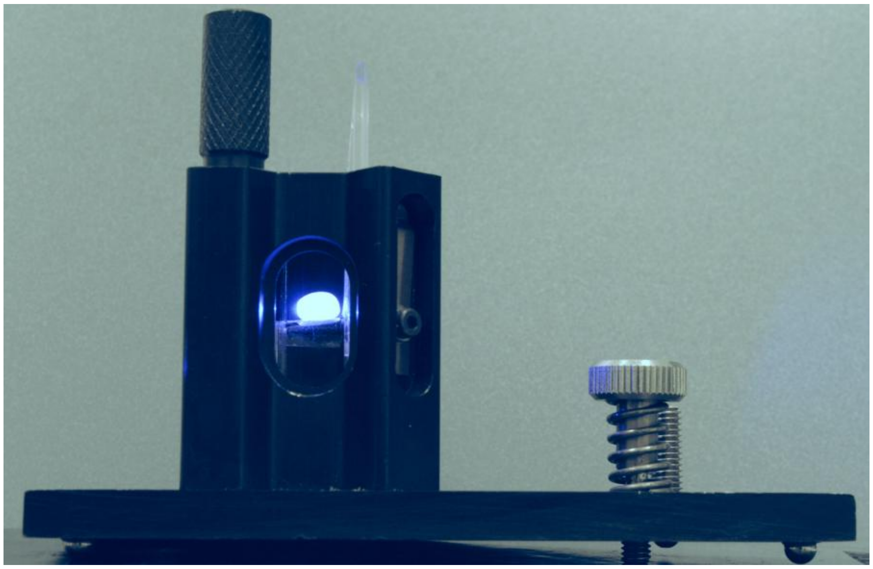

(b)

Fig S3. (a) Holder for millimeter-sized plastic pellets made from a polystyrene cuvette. (b) Plastic pellet mounted in the Varian Eclipse fluorimeter cuvette holder, illuminated with a 450 nm excitation beam.

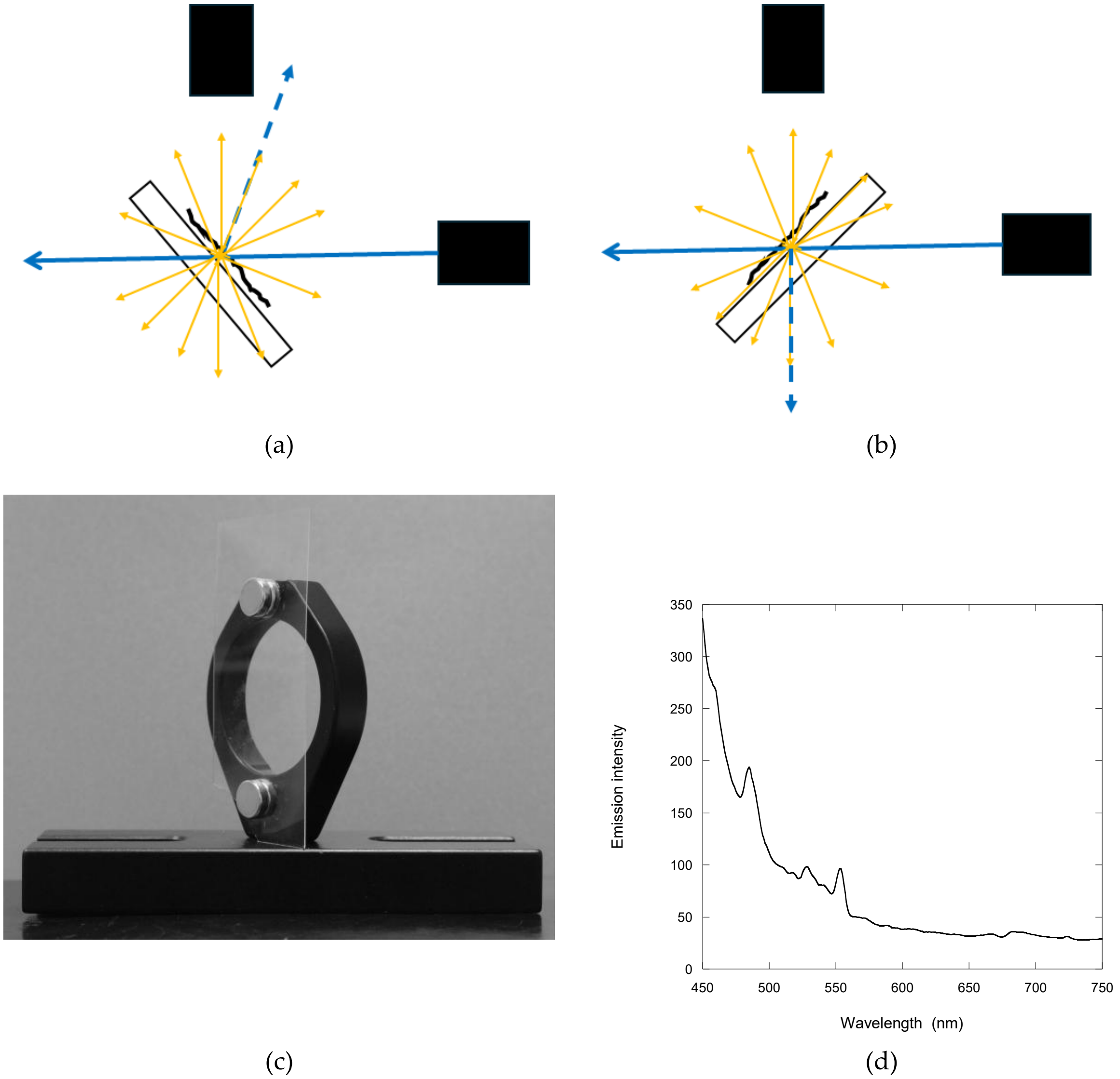


Fig S4. (a) Front-faced arrangement for holding plastic films adhered to a glass slide in a fluorimeter light path, angled at 40$^{o}$ to avoid direct reflection of the excitation beam (blue) into the emission detector. (b) Rear-faced arrangement to reflect the excitation beam away from the detector. (c) Holder for plastic films or samples dried on a glass slide for spectral analysis in a Varian Eclipse fluorimeter. (d) Scattered stray light detected by reflection of the excitation light (405 nm, 5 nm slit widths) from a glass slide mounted at ~ 40$^{o}$ to the excitation path in a Varian Eclipse fluorimeter showing "ghost" peaks at 485 nm, 529 nm and 552 nm.

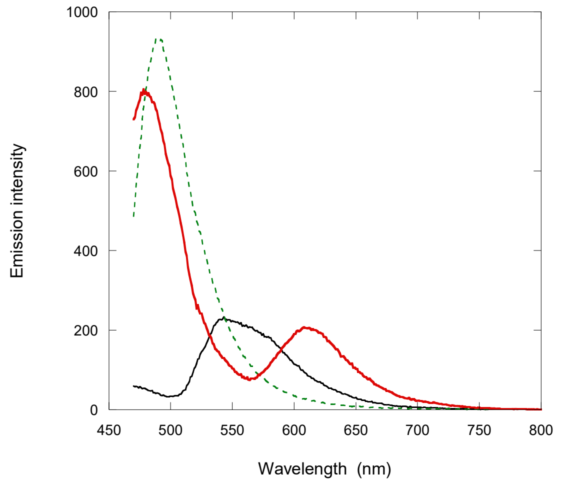


Fig S5. Emission spectra of DCVJ dissolved in isopropyl alcohol (green dashed), hexane (black) and a 70% hexane-30% IPA 1:1 mixture (red) showing peaks at 488 nm, 554 nm and 608 nm respectively.

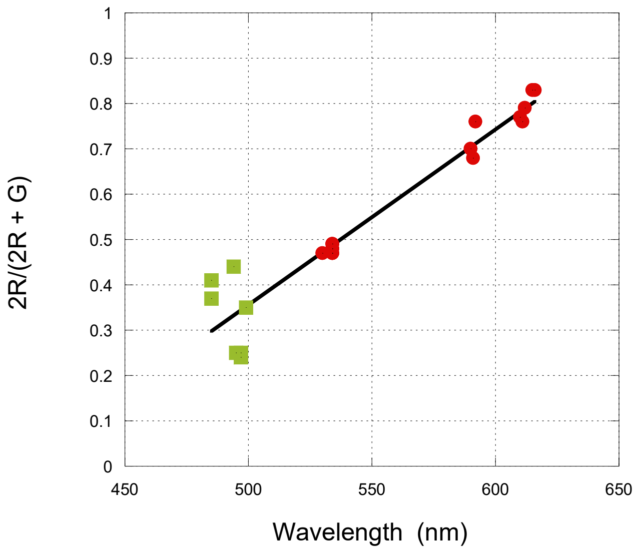


Fig S6. Correlation between the red chromaticity values (2R/(2R+G)) deduced from RGB values of Fig 3 and the peak emission wavelengths for Nile Red-stained (red circles) and DCVJ-stained (green squares) polymer samples determined by spectroscopy (Fig 4, Table 2). The chromaticity values in the low wavelength range are affected by the 515nm long pass filter (Fig S2) that significantly reduces the green component. Within the high wavelength range, the built-in IR of the Nikon D3000 reduces the red component (Fig S2a).

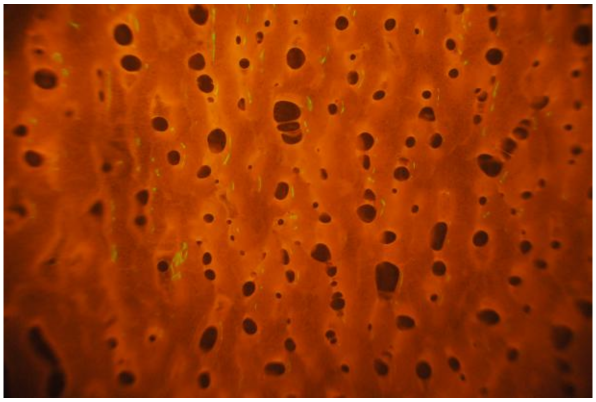

(a)

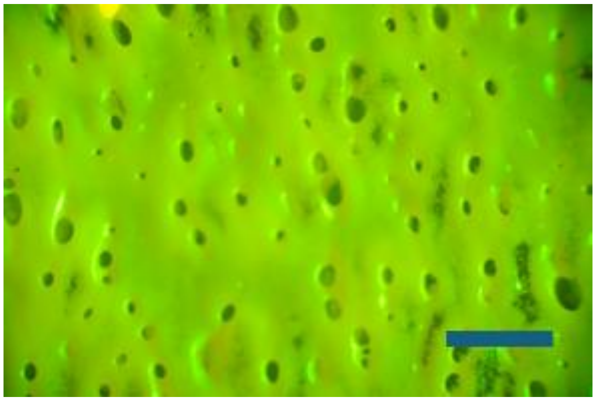

(b)

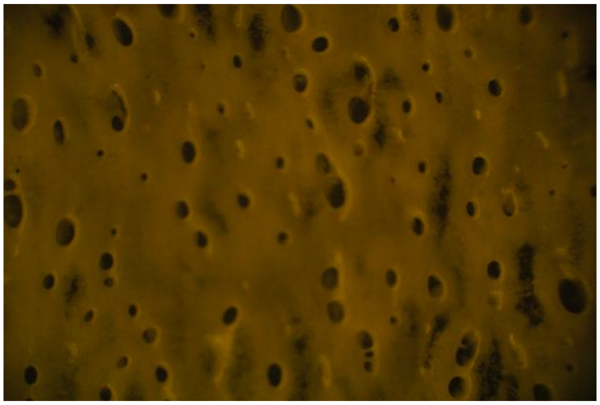

(c)

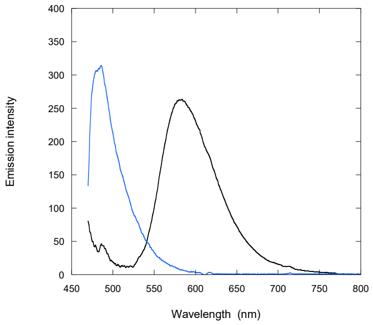


(d)

Fig S7. Staining of polystyrene Petri dishes in the presence of 98% acetone, 2% isopropyl alcohol. Scale bar = 1 mm in all panels. (a) Nile Red staining, $1/60^{th}$ s exposure, photographed through a 4x objective lens on an Amscope microscope with a Nikon D3000 camera and Zeiss LP515 nm filter. Chromaticity value, R/(R+G) = 0.72. (b) DCVJ staining with same set up and 1s exposure. Chromaticity value = 0.04. (c) Unstained control, treated with acetone and isopropyl alcohol only, with 1 s exposure. In the absence of acetone, Petri dishes were only weakly stained and lacked the roughened surface features. (d) Emission spectra recorded in front-faced mode (Fig S4a) for Nile Red (black) and DCVJ (blue) stained samples, corrected for the background scattering signal from the control sample.

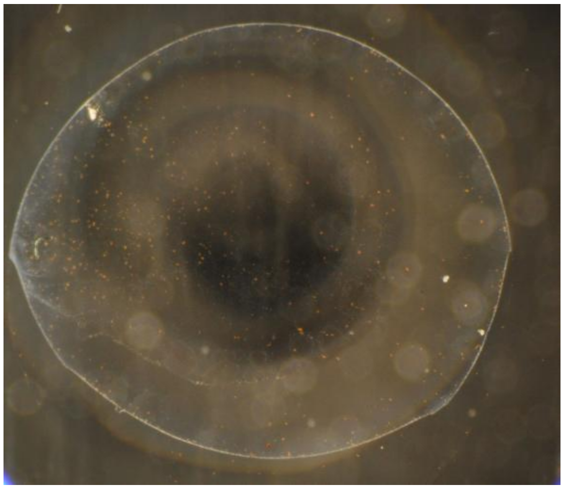

(a)

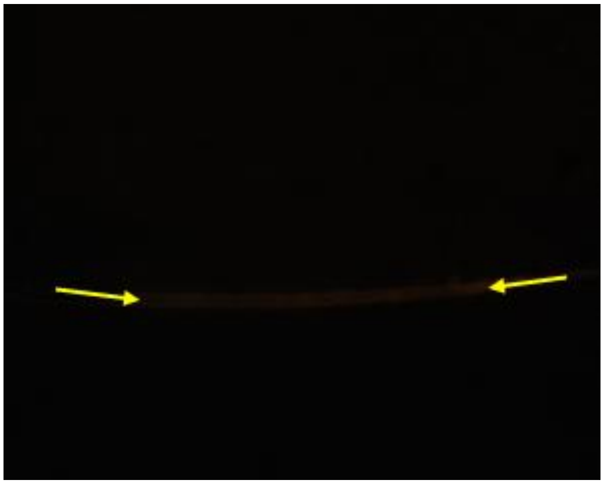

(b)

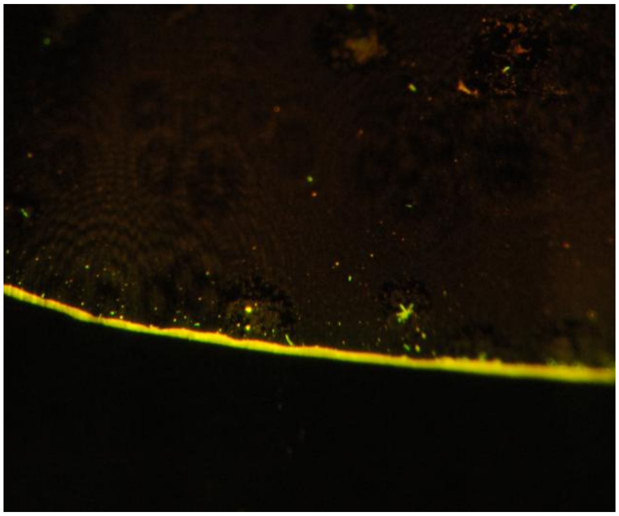

(c)

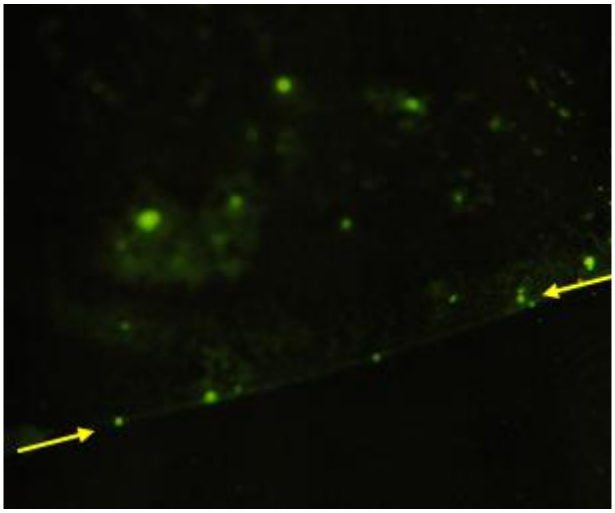

(d)

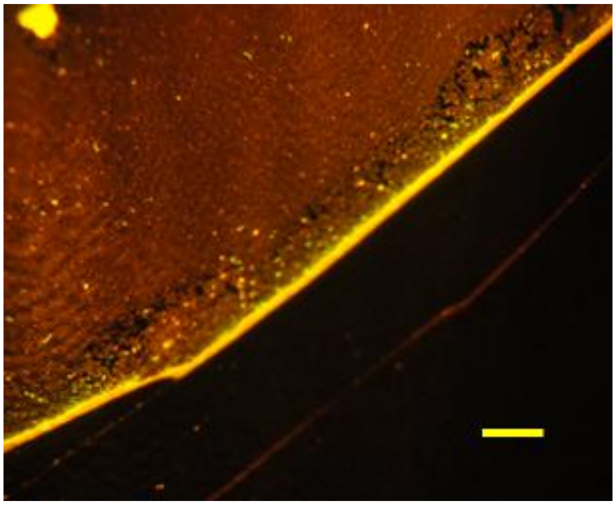

(e)

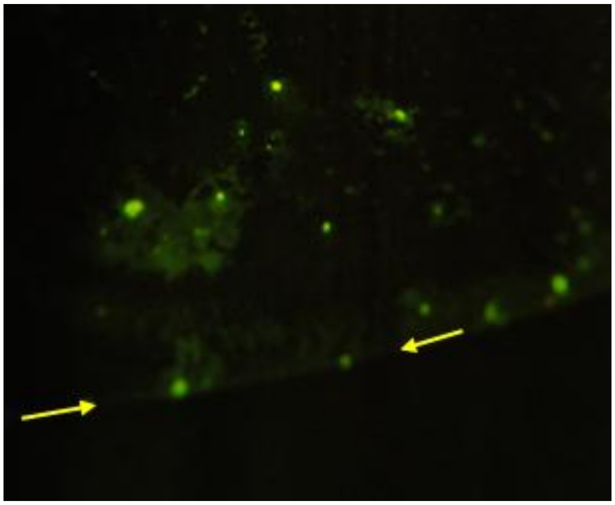

(f)

Fig S8. Stained 70 nm polystyrene beads (PS70) dried on slide and photographed with Nikon D3000 on an Amscope 120 microscope. (a) Darkfield view through a 2x objective lens of an evaporated 5 μL sample containing PS70 showing an outer ring (~4 mm diameter) of aggregated beads which scatter light. (b) to (f) Fluorescence images taken through a 20x 0.4 NA objective lens using a 450 nm laser for excitation and Zeiss 515nm long pass emission filter, 20 s exposure time. (b) Unstained PS70 beads showing low fluorescence background, but a weak boundary could be discerned from the breakthrough scattered light. (c) Edge of sample containing Nile Red-stained PS70 beads. (d) Sample containing evaporated Nile Red alone at the same concentration. (e) Edge of sample containing DCVJ-stained PS70 beads. (f) Sample containing evaporated DCVJ alone at the same concentration. Arrows show the location of the evaporation boundary in the absence of fluorescent beads. Analysis of raw images gave 2R/(2R+G) chromaticity values of NR = 0.78, NR+PS70 = 0.50, DCVJ = 0.82 and DCVJ+PS70 = 0.58 (compare with Fig 7c for the same solutions before evaporation). The scale bar in (e) = 10 μm and also applies to panels (b) to (f).
